# Two activation heat capacity regimes underlie temperature-dependent catalysis in homologous archaeal ADP-dependent kinases

**DOI:** 10.64898/2026.08.05.742859

**Authors:** Ignacio Aravena-Valenzuela, Pablo Maturana, Leslie Hernández-Cabello, Felipe Gonzalez-Ordenes, Victor Castro-Fernandez, Gabriel Vallejos-Baccelliere, Victoria Guixé

## Abstract

Enzyme activity increases with temperature up to a maximum, beyond which it declines, a behaviour traditionally attributed to thermal denaturation. However, some enzymes show activity decline well below the melting temperature. Macromolecular rate theory (MMRT) explains this phenomenon by introducing a negative activation heat capacity 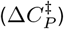, reflecting a transition-state ensemble more conformationally restricted than the ground state. Recently, 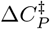 has been shown to be temperature-dependent and proposed as a general catalytic feature, though its variation within and across homologous families from distinct thermal niches remains unexplored. We characterized the glucokinase activity of three homologous bifunctional ADP-dependent PFK/GK enzymes: MbPFK/GK from the psychrotolerant *Methanococcoides burtonii*, MmPFK/GK from the mesophilic *Methanococcus maripaludis*, and ancM, the inferred ancestor of the *Methanococcales* order, which displays enhanced thermostability. MmPFK/GK and ancM display two 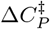 regimes, with abrupt changes in *k*_cat_ vs temperature: zero to moderately negative values at low temperatures, shifting sharply at elevated temperatures to highly negative values (−44 kJ mol^−1^ K^−1^ and −36 kJ mol^−1^ K^−1^, respectively), exceeding previous reports. Circular dichroism spectroscopy confirms that these extreme values reflect pre-melting conformational changes rather than denaturation. Despite being psychrotolerant, MbPFK/GK displayed the highest thermal stability 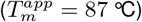 and a single 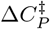 regime throughout all temperatures (−2.6 kJ mol^−1^ K^−1^). Domain-closure dynamics explain thermal adaptation and moderate-temperature 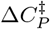 values; whereas the basis of the extreme high-temperature 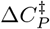 values remain unknown. To account for these two regimes, we present a two-pathway model incorporating a conformational equilibrium in which free enzyme and enzyme-substrate complex populate two catalytically competent conformations.

## Introduction

Through evolution, organisms have developed diverse adaptive strategies to maintain their metabolic activity across different thermal environments. Among these, a central mechanism is the optimisation of enzyme catalysis to ensure adequate reaction rates under extreme thermal conditions [1,2]. The Arrhenius and the Eyring–Polanyi theories both formalise this temperature dependence, predicting a linear relationship between the logarithm of the catalytic constant (*k*_cat_) and the reciprocal of temperature [3,4]. However, temperature often exerts opposing effects on catalysis and protein stability [5,6]. Enzyme activity typically increases with temperature up to a maximum, beyond which it declines. In general, departures from the Eyring-Polanyi linearity would be due to heat or cold-induced denaturation [1,5]. However, the existence of enzymes whose melting temperatures do not coincide with activity-decline temperatures challenges this view. Such phenomenon is increasingly recognised across a wide range of enzyme-catalysed reactions [7–18], and increasing evidence indicates that thermal denaturation alone is insufficient to fully account for it [7,11,17].

Macromolecular rate theory (MMRT) offers an alternative explanation for this phenomenon. It extends classical transition-state theory by incorporating temperature-dependent thermodynamic parameters through the introduction of an activation heat capacity parameter 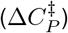. According to MMRT, a negative 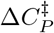 results in curvature in Arrhenius plots of enzyme-catalysed reactions, independent of protein denaturation, and thus clearly explains the decreases in activity observed well below the unfolding temperature [2]. The temperature dependence of *k*_cat_ ac-cording to MMRT is described by Eq. 1, where the terms 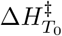 and 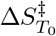 represent the activation enthalpy and entropy at a reference temperature, respectively.

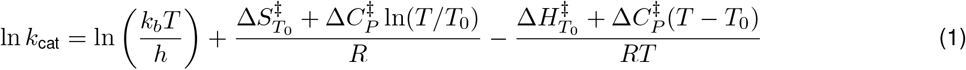

Over the past decade, negative 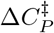 values have been reported for a growing number of enzymes, with typical magnitudes in the range of −1 to −12 kJ mol^−1^ K^−1^ [7,9,10]. Different interpretations have been given to this phenomenon. One of the most common is that it indicates a more conformationally restricted transition-state ensemble relative to the ground state [12,18–20]. Another example is that of computational studies on a psychrophilic *α*-amylase, which suggest that curvature can arise from the disruption of specific enzyme–substrate interactions at elevated temperatures rather than from protein unfolding, with the rate maximum occurring well below the melting temperature [21]. More recently, Walker et al. reported that 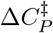 can be temperature-dependent, as they described in the glycosidase MalL, which exhibits a cooperative transition from an approximately zero value at low temperatures to −28.1 ± 6.3 kJ mol^−1^ K^−1^ at higher temperatures [17].

An interesting issue regarding how 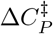 underlies thermoadaptation in enzyme homologs has recently been addressed. Nguyen et al. used ancestral sequence reconstruction of adenylate kinase to trace 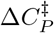 across the phylogeny, demonstrating that thermoadaptation involves a systematic redistribution of Δ*H*^*‡*^ and Δ*S*^*‡*^, with cold-adapted variants displaying lower Δ*H*^*‡*^ compensated by more negative Δ*S*^*‡*^ [11]. However, whether and how the temperature dependence of 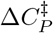 varies within individual enzymes across a homologous family occupying distinct thermal niches remains unexplored.

The ADP-dependent sugar kinases constitute a distinctive archaeal enzyme family that uses ADP rather than the canonical ATP as phosphate donor [22]. Members of this family exhibit either phosphofructokinase (PFK) activity or glucokinase (GK) activity, whereas others, including those from the methanogenic orders *Methanococcales* and *Methanosarcinales*, are bifunctional and catalyse both reactions at a single active site [23,24]. Since homologous enzymes have been characterised from organisms spanning a wide range of growth temperatures, this family provides an excellent model system for studying enzyme thermoadaptation [25]. Moreover, we have reconstructed the ancestral enzyme ancM, representing the last common ancestor of the enzymes of the *Methanococcales* order [26].

In this work, we use the MMRT framework to compare the temperature dependence of the glucokinase activity of three homologous bifunctional ADP-dependent PFK/GK enzymes spanning distinct thermal niches: MbPFK/GK from the psychrotolerant *Methanococcoides burtonii* (growth range 2.5–29 °C) [27], MmPFK/GK from the mesophilic *Methanococcus maripaludis* (optimal growth ~37 °C) [28], and ancM, the inferred ancestor of the *Methanococcales* order, which displays enhanced thermostability. We ask how 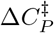 behaviour varies across this homologous family and whether it correlates with the thermal niche. The inclusion of ancM allows comparison with extant enzymes, bringing into the comparison a thermostable ancestral node within the same lineage, though the limited number of enzymes examined constrains the evolutionary conclusions that can be drawn. Moreover, in a previous report, we determined that conformational dynamics underlie the catalytic adaptability of this enzyme family across different temperatures [25], which is of great interest for addressing its relationship with 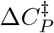 variation.

Our results show that MmPFK/GK and ancM exhibit an anomalous two-regime 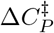 profile, involving an abrupt change in curvature in *k*_cat_ vs T curves, whereas MbPFK/GK shows a single regime. At moderate temperatures, the differences in 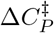 values among the three homologs correlate with the distinct conformational dynamics previously reported for this enzyme family [25]. Strikingly, at elevated temperatures, MmPFK/GK and ancM reach highly negative 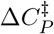 values that, to our knowledge, have not been previously reported for any enzyme. Importantly, these highly negative values occur in the absence of thermal denaturation. These findings expand the current framework for understanding the contribution of 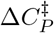 to catalytic thermoadaptation and raise new questions about the molecular mechanisms underlying such extreme activation heat capacity values.

## Results

### Temperature dependence of kinetic parameters

To characterize the temperature dependence of glucokinase activity, we determined glucose saturation curves across a range of temperatures for MbPFK/GK, MmPFK/GK, and ancM (Figure 1A–C; Table S1). The saturation curves for MmPFK/GK exhibited substrate inhibition at all temperatures assayed, with the inhibition constant (*K*_*I*_) increasing at temperatures below 20 °C and above 55 °C (Figure S1). This is an expected behaviour, since previous kinetic analysis has established that this inhibition arises from dead-end complex formation: glucose can bind non-productively to both the free enzyme before MgADP associates and to the enzyme-product complex before AMP dissociates [29]. In contrast, MbPFK/GK and ancM displayed classical Michaelis-Menten kinetics. The *K*_*m*_ values for glucose showed distinct temperature dependencies among the three enzymes: *K*_*m*_ decreased with temperature for MmPFK/GK, increased for MbPFK/GK, and remained relatively constant for ancM (Figure 1D). Notably, the increase in *K*_*m*_ with temperature for MbPFK/GK implies that this enzyme achieves its highest kinetic affinity for glucose at low temperatures, consistent with a cold-adaptation mechanism based on substrate binding optimization rather than rate enhancement [25].

**Figure 1:**
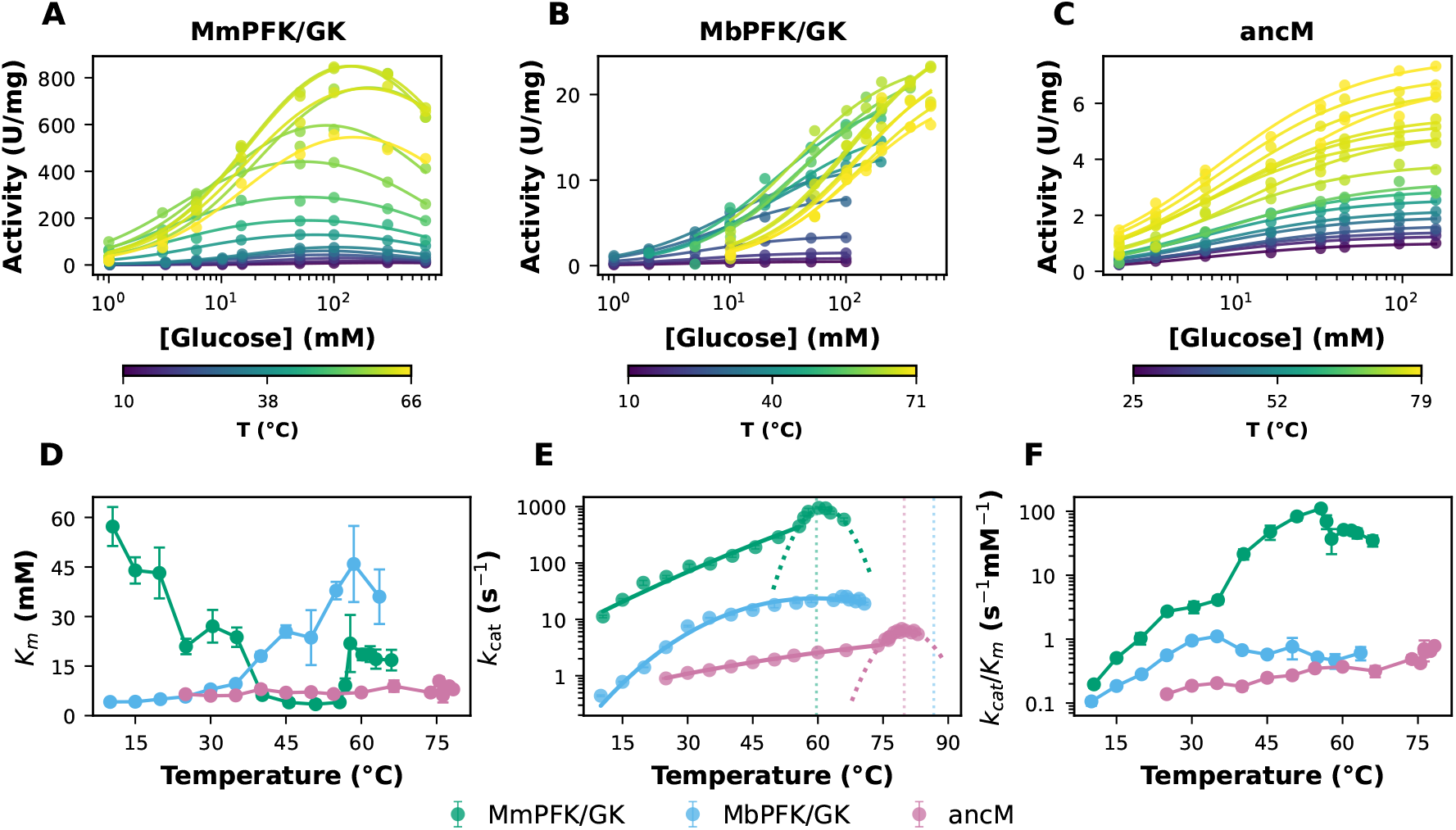
Temperature dependence of kinetic parameters for ADP-dependent PFK/GK enzymes. **(A–C)** Glucose saturation curves for MmPFK/GK (A), MbPFK/GK (B), and ancM (C); colour gradient from purple (lowest *T*) to yellow (highest *T*). Solid lines represent the fitting of the Michaelis–Menten (Eq. 8) or substrate inhibition model (Eq. 9, panel A). **(D)** *K*_*m*_ for glucose, **(E)** *k*_cat_, and **(F)** *k*_cat_*/K*_*m*_ vs temperature for MmPFK/GK (green), MbPFK/GK (blue), and ancM (pink). Solid lines in (E) are Eyring fit below 55.7 °C for MmPFK/GK and below 73.8 °C for ancM, while MMRT equation was fitted above those temperatures (dashed lines, Eq. 1; Table 1). In (E), the transition from solid to dashed lines marks the boundary between the low/moderate-temperature and high-temperature 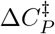 regimes, occurring at 55.7 °C for MmPFK/GK and 73.8 °C for ancM. Vertical dotted lines in (E) mark each enzyme’s 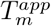. Data are mean ± fit SEM.

The *k*_cat_ values increased with temperature for all three enzymes, reaching a maximum, then declined at higher temperatures (Figure 1E). The temperature of maximal *k*_cat_ differed among enzymes: approximately 60 °C for MmPFK/GK, 66 °C for MbPFK/GK, and 79 °C for ancM (Table S1). The *k*_cat_ values varied considerably, with MmPFK/GK displaying the highest values (up to ~ 950 s^−1^), followed by MbPFK/GK (~25 s^−1^) and ancM (~7 s^−1^). The catalytic efficiency (*k*_cat_*/K*_*m*_) reached maximal values at approximately 55 °C for MmPFK/GK and 35 °C for MbPFK/GK, whereas for ancM the catalytic efficiency continued to increase throughout the assayed temperature range (Figure 1F).

**Table 1:**
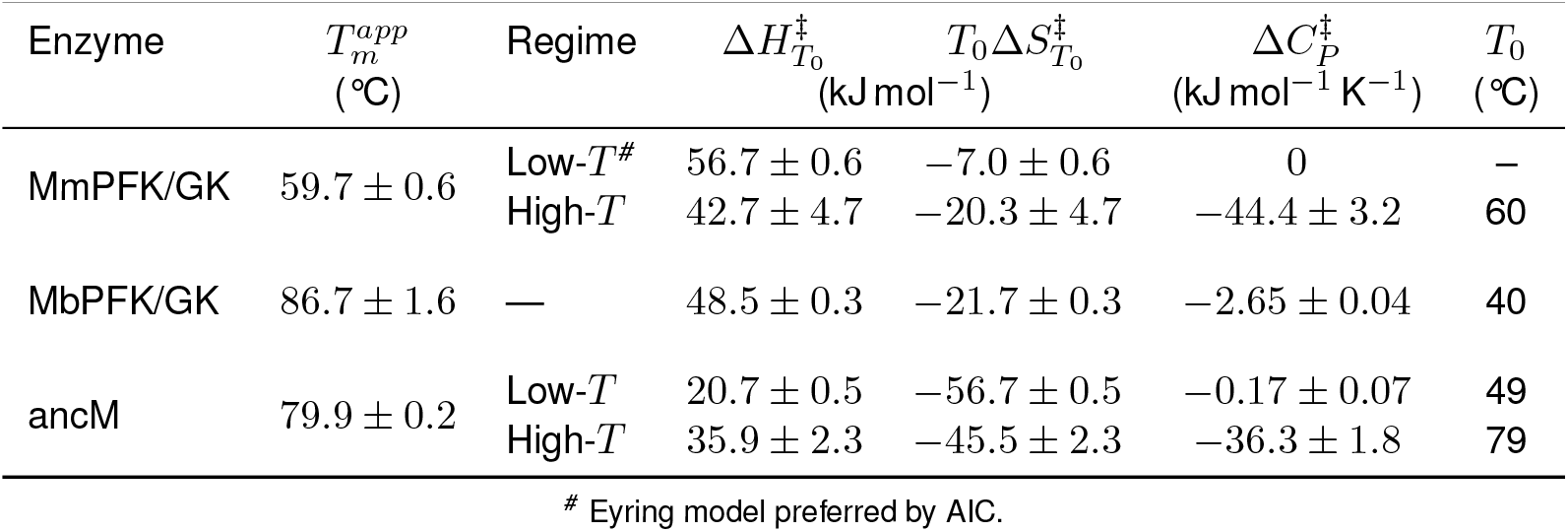
Activation parameters and thermal stability. Kinetic parameters were obtained by fitting *k*_cat_ to the Eyring^*#*^ or MMRT model.

Notably, the *k*_cat_ curves for MmPFK/GK and ancM exhibit a pronounced change in curvature at elevated temperatures (Figure 1E), clearly showing two kinetically distinct regions (solid lines and dashed lines). This two-regime behaviour is analysed in terms of activation heat capacity in the following section. In Figure 1E, vertical dotted lines mark each enzyme’s 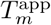.

### Activation heat capacity profiles

To analyse the temperature dependence of catalytic activity within the MMRT framework, we analyse the *k*_cat_ vs *T* data in terms of MMRT or Eyring models. The analysis revealed distinct 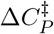 profiles among the three enzymes (Figure 1E, Table 1). MbPFK/GK was well described by a single-regime MMRT fit across the entire temperature range, whereas MmPFK/GK and ancM exhibit curvature changes at high temperatures, with two recognizable regimes, indicating a transition in the thermodynamic properties governing catalysis. In the first stage of our analysis, each regime was treated independently, separating a low-to-moderate temperature regime (below 55.7 °C for MmPFK/GK and below 73.8 °C for ancM), from a high temperature regime above these thresholds. The temperature dependence of the individual thermodynamic contributions computed from the fitted MMRT parameters is shown in Table 1 and Figure S3.

In the moderate temperature regime, AIC-based comparison between the classical Eyring equation 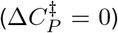 and the MMRT model (Figure S2, Table S5) identified the Eyring model as the best fit for MmPFK/GK, whereas the MMRT model was preferred for ancM. For MmPFK/GK, the estimated 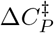 was 0 kJ mol^−1^ K^−1^, indicating that it was statistically indistinguishable from zero in this regime. In contrast, for ancM the fitted MMRT model exhibits a small but significant negative 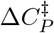, with a fitted value of − 0.17 ±0.07 kJ mol^−1^ K^−1^. These moderate-regime values, together with the single-regime value for MbPFK/GK, fall within the range typically reported for other enzymes [7,9,10].

At elevated temperatures, MMRT equation was fitted to the MmPFK/GK and ancM *k*_cat_ vs *T* curves. This resulted in highly negative 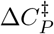 values: −44.4 ± 3.2 kJ mol^−1^ K^−1^ for MmPFK/GK and −36.3 ± 1.8 kJ mol^−1^ K^−1^ for ancM. Although two-regime 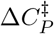 behaviour has been recently described for other enzymes [17], the magnitudes observed here far exceed previously reported values. According to the main interpretations of the MMRT framework, such extreme 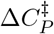 values would correspond to a dramatic narrowing of the conformational ensemble accessible to the transition state relative to the ground state [12,18–20]. This highly contrasts with MbPFK/GK, which, besides not showing a curvature transition, showed a 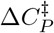 of −2.65 ± 0.04 kJ mol^−1^ K^−1^, that is within the typical range reported [7,17].

Experimental controls were performed for the auxiliary coupling enzymes used for the kinetic assays. Auxiliary enzymes display increasing activity across their respective operational ranges, with no curvature or thermal anomaly that could be projected onto the ADP-PFK/GK *k*_cat_ vs temperature data: LmG6PDH (from the mesophilic *Leuconostoc mesenteroides*, used at 10–45 °C; Figure S5, Table S4) and TmG6PDH (from the hyperthermophile *Thermotoga maritima*, used at 50–83 °C; Table S3). On the other hand, the temperatures of regime transition differ between MmPFK/GK and ancM in parallel with their respective 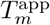 values, thereby excluding any artifact arising from the coupled assay used to measure enzyme activity. Therefore, the two-regime 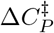 profiles are intrinsic to MmPFK/GK and ancM.

The thermodynamic decomposition of the fitted parameters (Figure S3) shows that in the high-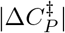 regimes, Δ*H*^*‡*^ and *T* Δ*S*^*‡*^ change steeply and in parallel while Δ*G*^*‡*^ varies only modestly, consistent with enthalpy–entropy compensation.

### Thermal stability and secondary structure changes

To assess whether the decline in *k*_cat_ at elevated temperatures could be attributed to thermal denaturation, we monitored protein unfolding by CD spectroscopy (Figure 2A, Table 1). The transition to the highly negative 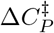 regime coincides with proximity to 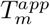 for both MmPFK/GK and ancM.

**Figure 2:**
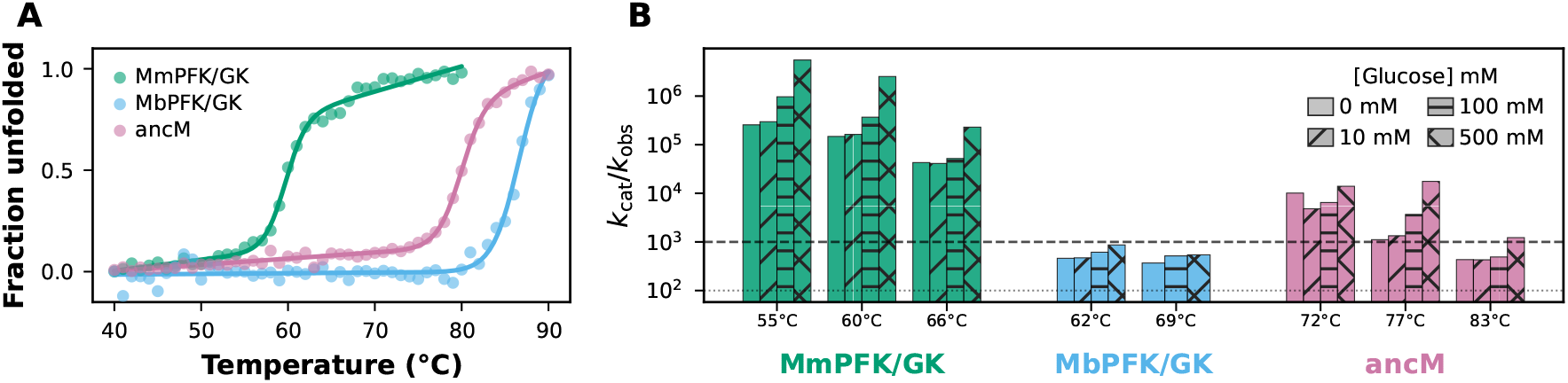
Thermal stability and unfolding kinetics. **(A)** Thermal unfolding monitored by CD at 222 nm for MmPFK/GK (green), MbPFK/GK (blue), and ancM (pink); solid lines correspond to the fitting of Eq. 10. **(B)** *k*_cat_*/k*_obs_ ratios near the onset of thermal unfolding; bars coloured by enzyme and hatched by glucose concentration (see legend). Horizontal lines mark ratios of 10^2^ (dotted) and 10^3^ (dashed).

A critical question is whether *k*_cat_ values at these elevated temperatures genuinely reflect catalysis by the native enzyme. To address this, we monitored time-resolved unfolding kinetics by CD at 222 nm at the same temperatures used in the kinetic assays, in the presence of different glucose concentrations (0–500 mM, Figure S6).

*k*_obs_ is the first-order rate constant describing the time-dependent loss of native secondary structure, obtained by fitting the CD signal at 222 nm over time to Eq. 11. Because *k*_cat_ and *k*_obs_ are both first-order rate constants determined under matched conditions, their ratio directly indicates how many catalytic turnovers occur, on average, before an enzyme molecule loses its native fold. The resulting *k*_cat_*/k*_obs_ ratios are shown in Figure 2B.

Comparison of CD time-course amplitudes with equilibrium spectra revealed that *k*_obs_ describes enzyme-specific processes. This unfolding is irreversible under our conditions. For MmPFK/GK at 60–66 °C, the CD signal approaches fully denatured values, consistent with complete unfolding 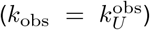. In contrast, for ancM and MbPFK/GK the signal plateaus at intermediate levels, indicating a transition to a defined partially unfolded intermediate 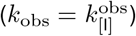. These two scenarios are described by:

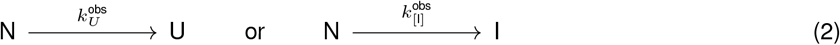

where N, U, and I denote the native, fully unfolded, and partially unfolded intermediate states, respectively.

For MmPFK/GK,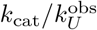 ratios exceeded 10^4^ throughout the assayed range, and the time-course data revealed a lag phase preceding cooperative unfolding (Figure S6A, D). For ancM, 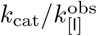 remained above 10^3^ except at 83 °C, where it decreased to a few hundred (Figure 2B), and the CD signal plateaued at a level consistent with a defined partially unfolded intermediate (Figure S6H). For MbPFK/GK, ratios were of order 10^2^–10^3^ at 62–69 °C, with the CD time course also plateauing at a fraction unfolded of ~ 0.2 at 62 °C (Figure S6F, I). The three enzymes were stabilized as glucose concentration increased (Figure S7); at 500 mM, *k*_obs_ was reduced by factors ranging from 5 to 22 for MmPFK/GK, 3 to 16 for ancM, and 1.5 to 2 for MbPFK/GK relative to the glucose-free condition, depending on temperature.

Together, these data show that catalytic turnover precedes unfolding, even at elevated temperatures. Thus, initial velocity measurements were performed before significant unfolding occurred. The *k*_cat_ values in this regime, therefore, reflect enzyme species that retain catalytic competence despite incipient conformational destabilization, rather than a post-denaturation artifact.

To investigate the structural changes accompanying increasing temperature, we recorded far-UV CD spectra at different temperatures in the absence and presence of 100 mM glucose (Figure 3A). At 25 °C, all three enzymes exhibited the characteristic α-helical CD signature, with minima at approximately 208 and 222 nm (Figure 3B).

**Figure 3:**
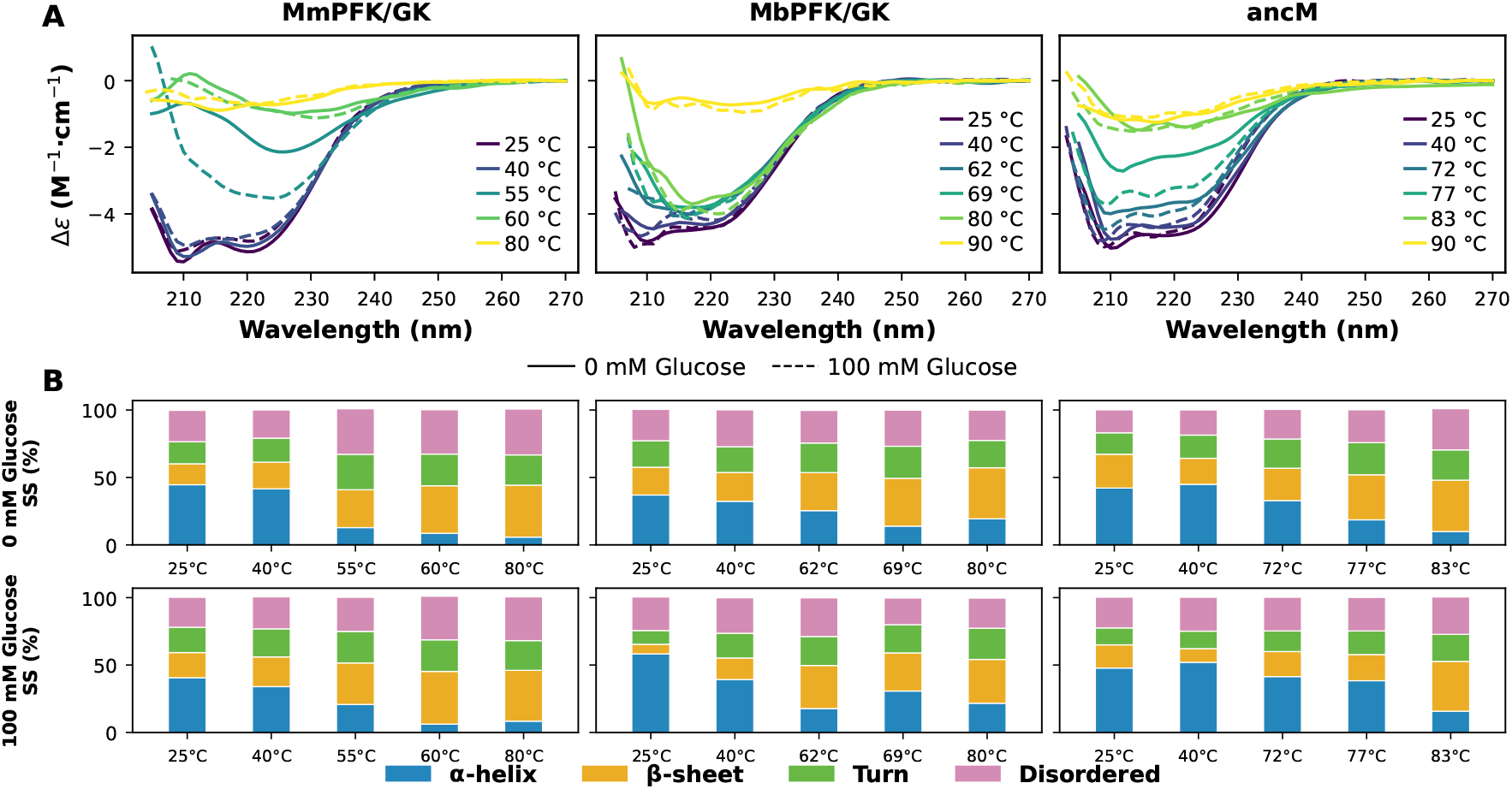
Far-UV CD spectra and secondary structure content. **(A)** CD spectra at selected temperatures for MmPFK/GK (left), MbPFK/GK (centre), and ancM (right); solid lines, no glucose; dashed, 100 mM glucose; colour gradient from purple to yellow indicates increasing temperature. **(B)** Secondary structure percentages (*α*-helix, *β*-sheet, turn, disordered). Columns: enzymes; rows: 0 mM (top) and 100 mM glucose (bottom).

At elevated temperatures, the CD spectra analysis revealed enzyme-specific losses of *α*-helical content accompanied by reciprocal gains in *β*-sheet fraction (Figure 3B). Without glucose, the transition was abrupt for MmPFK/GK, with *α*-helix dropping from ~38% at 25 °C to ~10% at 60 °C 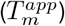, with *β*-sheet rising correspondingly. For MbPFK/GK the loss was more gradual, retaining ~17% *α*-helix at ~62 °C and *β*-sheet increasing to~34%. ancM showed the most pronounced glucose-dependent structural protection: in the presence of 100 mM glucose, *α*-helix content was ~41% at 25–40 °C, declining gradually to ~35% at 72–77 °C before dropping to ~16% at 83 °C, whereas in the absence of glucose the loss of *α*-helical content began at lower temperatures, reaching ~18% already at 77 °C. This glucose-dependent structural protection is qualitatively concordant with the kinetic stabilization observed in the time-resolved CD experiments.

### Domain-closure dynamics correlate with 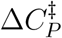

The distinct conformational dynamics of MbPFK/GK and MmPFK/GK provide an opportunity to test whether the mode of domain-closure motion influences 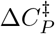. Previous work established that MbPFK/GK undergoes “breathing-type” domain motion whereas MmPFK/GK exhibits “twist-type” motion, and that a quadruple mutant M4MbPFK/GK (A336K/S339D/N110E/V463K), in which two salt bridges from the mesophilic branch were introduced into the psychrophilic scaffold, partially shifts domain motion from breathing-toward twist-type [25].

We performed a kinetic characterization of M4MbPFK/GK across 10–60.5 °C (Figure S8). Its *k*_cat_ values were lower than those of MbPFK/GK across the full temperature range, and the MMRT model was preferred over the Eyring model by AIC (Figure 4A and Figure S8B; Table S7), yielding 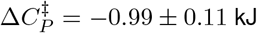 mol^−1^ K^−1^ (Figure 4B). The *K*_*m*_ for glucose increased steeply above 45 °C (Figure S8C), and *k*_cat_*/K*_*m*_ was lower than that of MbPFK/GK at most temperatures (Figure S8D). No second 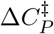 regime was observed within the measured range.

**Figure 4:**
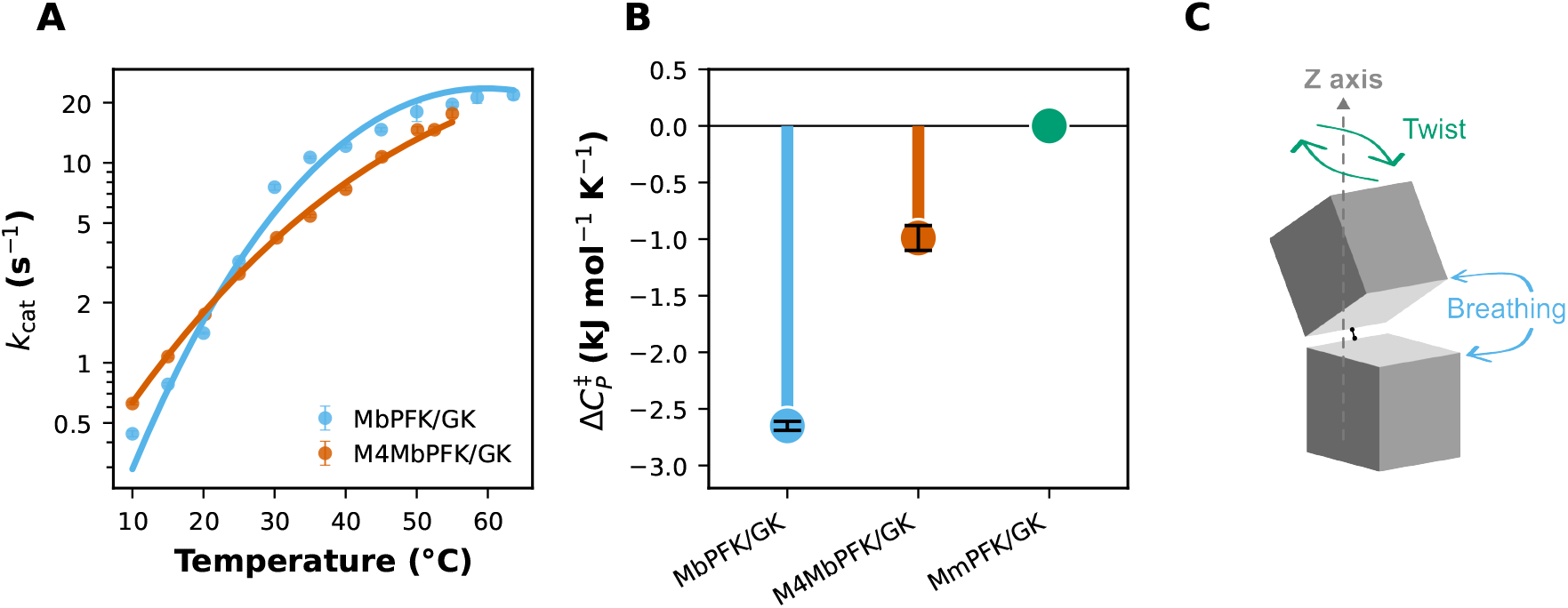
Domain-closure dynamics and 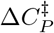. **(A)** *k*_cat_ vs temperature for MbPFK/GK (blue) and M4MbPFK/GK (orange); lines, MMRT fits; mean ± SEM. **(B)** Moderate-temperature 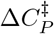 ordered by domain-closure mode and *k*_cat_*/K*_*m*_. **(C)** Schematic of the two domain-closure modes: breathing motion along the interdomain (Z) axis (blue) and twist motion off the axis (green).

The two closure modes differ in their underlying motion (Figure 4C): in the psychrophilic MbPFK/GK, the domains move along the interdomain (Z) axis in a breathing motion, directly approaching and separating, whereas in the mesophilic MmPFK/GK closure occurs through a more pronounced twist motion off the Z axis, conferred by the two salt bridges absent in the psychrophilic lineage, with M4MbPFK/GK adopting twist-type dynamics intermediate between the two proteins. Arranging the moderate-temperature 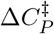 values by domain-closure mode reveals a consistent trend: MbPFK/GK (breathing, −2.65 kJ mol^−1^ K^−1^) > M4MbPFK/GK (intermediate, −0.99 kJ mol^−1^ K^−1^) > MmPFK/GK (twist,≈ 0; Eyring preferred) (Figure 4B). This arrangement supports the view that the mode of domain-closure motion shapes the degree of conformational restriction imposed during transition-state formation.

## Discussion

It has been proposed that a temperature-dependent 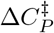 is a general feature of enzyme catalysis, prompting its examination in additional enzymatic systems [17]. Our results confirm that this behaviour extends to homologous ADP-dependent PFK/GK enzymes. However, in our case, the observed temperature dependence diverges from previously reported patterns in the literature. This includes a pronounced change in curvature, as seen in MmPFK/GK and ancM, and also 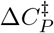 magnitudes in the high-temperature regime (− 44 and − 36 kJ mol^−1^ K^−1^ for MmPFK/GK and ancM, respectively) that far exceed any previously reported values, including the 20.8 kJ mol^−1^ K^−1^ obtained by Walker et al. from independent segment fits to the high-temperature data of MalL [17].

### Methodological concerns regarding the determination of 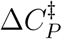

Several methodological considerations concern the experimental measurement of 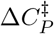 and hence possible errors in its interpretation. The MMRT framework strictly applies to the chemical step of the catalysed reaction, accounted for by *k*_cat_; if activity is measured at sub-saturating substrate concentration, then the temperature effect on substrate binding will also affect the dependence of initial velocity on temperature, so the resulting 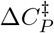 becomes apparent rather than true 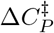. We illustrate this by analysing the initial velocity vs temperature data obtained at different glucose concentrations for MmPFK/GK and fitting MMRT in the high-temperature regime (Figure 5A). The estimated 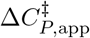 values vary four-fold across this range, from − 13 to − 50 kJ mol^−1^ K^−1^ (Figure 5B), as glucose concentration varies. Many published MMRT analyses do not document the saturation state across the temperature range assayed. For example, studies that compile rate–temperature datasets from multiple publications, such as Arcus and Pudney [8], do not explicitly state whether the reported parameters were derived from measurements performed under substrate-saturating conditions.

**Figure 5:**
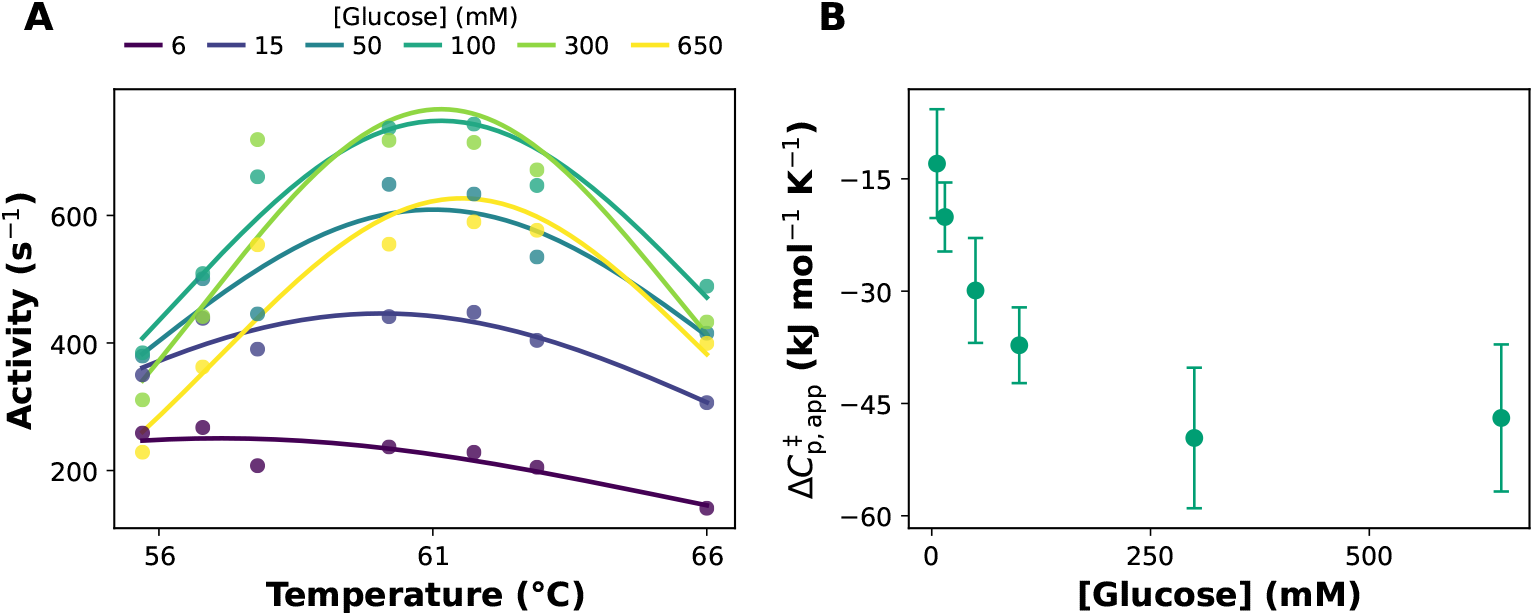
Apparent 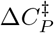 dependence on substrate saturation. **(A)** MmPFK/GK activity expressed in s^−1^ vs temperature in the high-temperature regime at six glucose concentrations (legend, mM); lines correspond to independent MMRT fits per concentration. **(B)** Fitted apparent 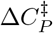 vs glucose concentration; error bars correspond to the standard error of the MMRT fit.

A related ambiguity affects analyses that fit MMRT to *k*_cat_*/K*_*m*_: the parameter bundles binding with the chemical step, so the extracted 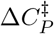 may largely reflect a heat capacity change of binding 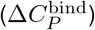 rather than activation, as noted by Åqvist [16]. Bunzel et al. adopted the *k*_cat_*/K*_*m*_ regime for evolved Kemp eliminases due to limited substrate solubility, arguing that the much larger evolutionary change in *k*_cat_ relative to *K*_*m*_ supports a chemical interpretation [14]; the binding contribution is nonetheless not quantitatively separated. A converse issue arises when substrate inhibition is overlooked. For example, the cold-adapted lipase data of van der Ent et al. [30] exhibit velocity curves consistent with substrate inhibition, yet the canonical Michaelis–Menten model was used for fitting, likely biasing the estimated *k*_cat_. In our case study, MmPFK/GK itself shows glucose substrate inhibition at all temperatures (Figure 1A), which we explicitly account for using the substrate inhibition model (Eq. 9).

### Secondary-structure changes reveal partial rather than complete unfolding

CD spectra decomposition reveals a progressive loss of α-helical content with reciprocal accumulation of *β*-sheet (Figure 3B), further supporting partial rather than complete unfolding. This pattern is reminiscent of the partial destabilization observed for psychrophilic α-amylase [21], where disruption of specific interactions rather than complete unfolding underlies the temperature maximum. Hence, it would be expected that temperature-induced conformational changes could be a relevant factor in the temperature dependence of catalysis. An important caveat is that thermal unfolding under these conditions is irreversible, so the population of catalytically competent enzyme decreases over time. However, the *k*_cat_*/k*_obs_ ratios exceeding 10^3^ mean that >99.9% of catalytic turnovers occur before appreciable loss of activity, and *k*_cat_ is determined from initial velocities of a nearly homogeneous native population. Nevertheless, the MMRT parameters in the high-temperature regime should be regarded as apparent values whose physical interpretation may be complicated by population-level effects from incipient destabilization of the ground-state ensemble, opening the possibility that more than one catalytically active conformation is populated within the ground-state ensemble.

### Analysis of the dual-regime 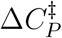 behaviour under the TLC framework

Enzymatic catalysis is inherently a multi-step process, in which each elementary step possesses its own Δ*H*^*‡*^ and Δ*S*^*‡*^ governed by independent heat-capacity terms [9]. The observed dual-regime behaviour departs from previous descriptions, suggesting that temperature differentially affects the individual steps of the catalytic cycle.

While the thermodynamic relationship linking Δ*H*^*‡*^ and Δ*S*^*‡*^ through 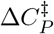 holds for each elementary step individually, this relationship need not hold for the apparent activation parameters obtained by fitting experimental data with a single-barrier model such as Eq. 1. Our first approach was therefore a semi-empirical analysis in which the *k*_cat_ vs *T* curve is divided into two segments according to the change in curvature, fitting a separate function to each and leaving aside theoretical considerations.

A plausible way to account for the dual behaviour detected in this work is to consider the differential effect of temperature on the intermediate steps occurring between substrate binding and product release. This could be done, for example, by separating the chemical step from product release within the Michaelis–Menten scheme, E + S *⇌* ES *⇌* EP *→* E + P; or by introducing a conformational change, E + S *⇌* ES *⇌* E^*′*^S *→* E + P. Because each step is differentially affected by temperature, two-regime behaviour may arise from temperature-dependent changes in the rate-limiting step of the catalytic cycle.

A conceptual framework of this type was proposed by Walker et al. [17], who attributed the dual behaviour described in their example to an intermediate step in the catalytic sequence, in which the enzyme–substrate complex (ES) is in equilibrium with a “transition-state-like conformation” (TLC) that preferentially promotes the chemical step. The cooperative nature of this conformational transition gives rise to a sigmoidal temperature dependence of 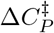, which is described by Eq. 3:

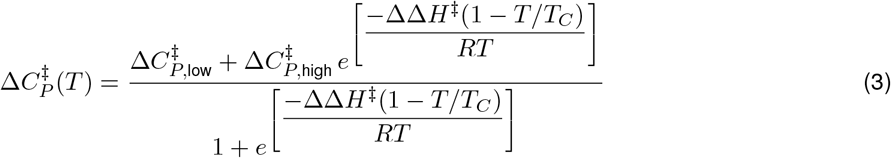

where *T*_*C*_ is the midpoint of the conformational transition, 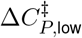 and 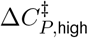 are the limiting activation heat capacities in each regime, and ΔΔ*H*^*‡*^ governs the cooperativity of the transition. For data sets with fewer temperature points, they also proposed a linear approximation (MMRT-1L, Eq. 4):

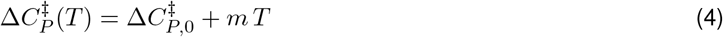

where 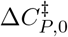 is an intercept and *m* is the slope, capturing both the magnitude and steepness of the 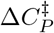 transition in a single additional parameter relative to MMRT. In the MMRT-2S model, a negative 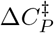 arises from the reduced conformational ensemble of the TLC relative to the ES complex; a broadening of this ensemble difference would amplify the heat capacity difference between ground and transition states, potentially accounting for the extreme 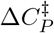 magnitudes observed here.

To assess whether the curvature change observed at high temperatures in MmPFK/GK and ancM is explained by a change in the rate-limiting step of the catalytic cycle—either through separation of the chemical step from product release or through the formation of an intermediate such as a TLC—we analysed our data following a general three-step model. Assuming steady-state and deriving the Michaelis–Menten equation, the following expression for *k*_cat_ is obtained as a function of the first-order microscopic rate constants for the steps after substrate binding:

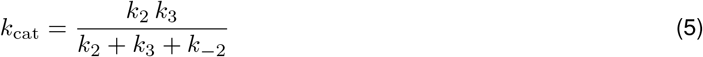

The microscopic constants *k*_2_ and *k*_3_ correspond to the two first-order steps occurring after substrate binding. Adding intermediate steps and accounting for the temperature dependence of each is inherently complex, as it multiplies the number of model parameters, especially when MMRT behaviour is assigned to more than one rate constant. To address this issue, Walker et al. derived a simplified fitting equation based on a series of approximations for fitting the empirical data [17]. In our case, even without such simplifications, the three-step model proved impossible to fit. This outcome is mathematically expected: the steady-state *k*_cat_ depends hyperbolically on each of the first-order microscopic constants *k*_2_ and *k*_3_ (Eq. 5), a relationship that holds regardless of how many intermediate steps are considered. As a result, a temperature-induced shift in the rate-limiting step produces a smooth, monotonic change in apparent thermodynamic parameters rather than an abrupt transition with a positive change in slope toward extreme 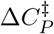. The thermodynamic parameterizations derived from it, MMRT-1L (Eq. 4) and MMRT-2S (Eq. 3) [17], are likewise rejected by AICc, with fit curves indistinguishable from those of standard MMRT (Figure S9, Table S8).

### A two-pathway model accounts for the dual-regime 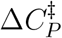 behaviour across the entire temperature range

The preceding analysis shows that a linear catalytic pathway does not account for our experimental data. We propose instead a two-pathway model incorporating a conformational equilibrium in which both the free enzyme and its enzyme–substrate complex populate two catalytically competent conformations, E/E^*′*^ and ES/E^*′*^S:

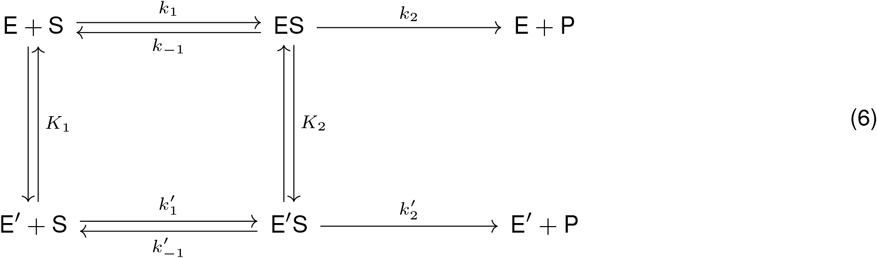

where *K*_1_ and *K*_2_ are the conformational equilibria of the free enzyme and the enzyme–substrate complex, respectively, and 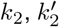 are the catalytic rate constants from ES and E^′^S. Under rapid equilibrium and assuming equivalent substrate affinities for E and E^′^, *k*_cat_ is a *K*_2_-weighted average of the two rates:

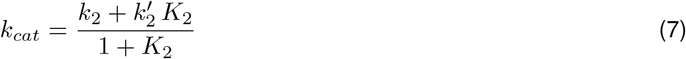

At low temperatures ES dominates (*K*_2_ *≪* 1) and *k*_cat_ *≈ k*_2_; as temperature rises near 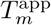 the population of E^*′*^S grows and *k*_cat_ shifts toward 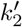. If *k*_2_ follows Eyring and 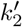 follows MMRT with a highly negative 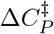, then the *K*_2_-weighting amplifies the MMRT contribution cooperatively, generating the rate maximum and the abrupt 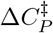 transition without assuming a discrete regime boundary.

The model was successfully fitted to *k*_cat_ vs *T* for both MmPFK/GK and ancM (Figure 6), with *K*_2_ treated as temperature-independent and fitting restricted to *k*_cat_ data since high-temperature assay conditions can introduce additional uncertainty in *K*_*m*_. Fit accuracy is comparable to the piecewise MMRT fitting used as a reference. Parameter determination remains challenging due to inherent correlations, as Walker et al. noted for the MMRT-2S model [17].

**Figure 6:**
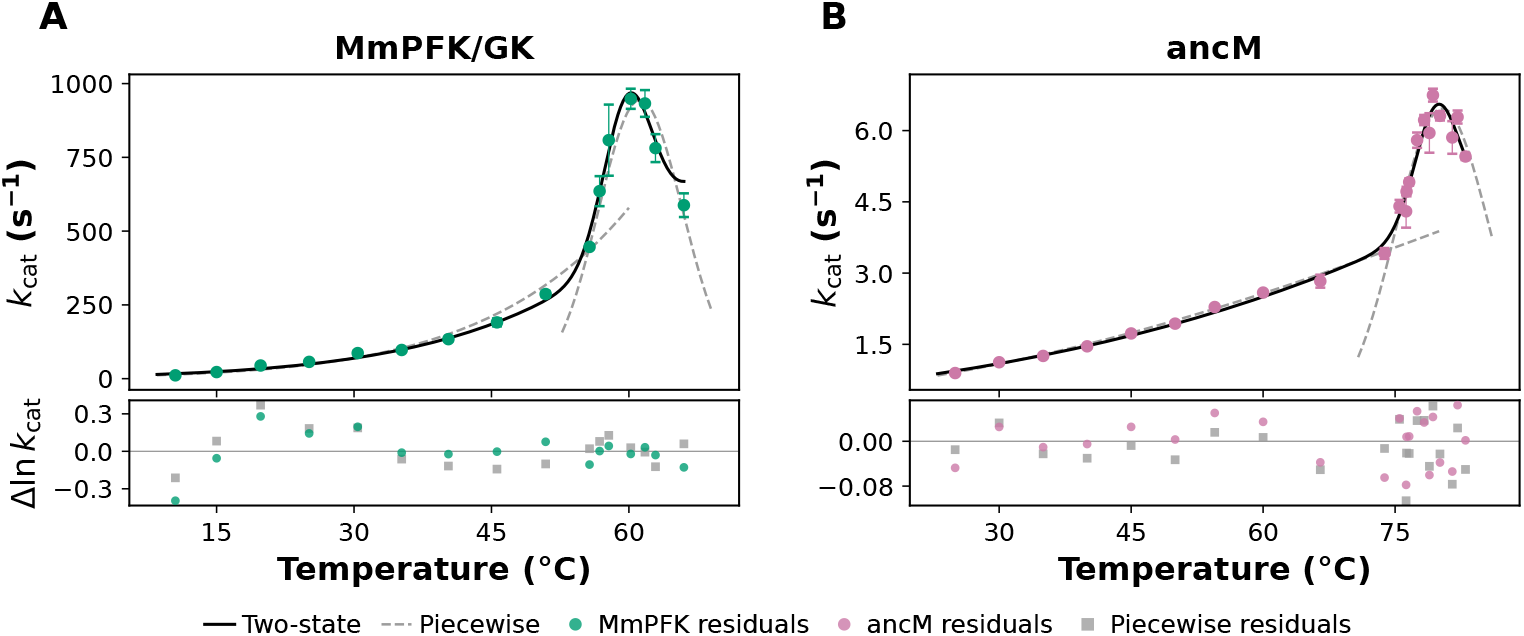
Two-state conformational model (Eq. 7) fitted to *k*_cat_ for MmPFK/GK and ancM, with *k*_2_ following Eyring, 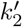 following MMRT, and *K*_2_ temperature-independent. **(A)** MmPFK/GK (RMSE 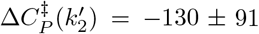 kJ mol^−1^ K^−1^). **(B)** ancM 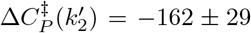 kJ mol^−1^ K^−1^). Dashed lines: piecewise reference (Eyring+MMRT for MmPFK/GK, RMSE = 0.148; two-MMRT for ancM, RMSE = 0.042). Lower panels: residuals Δ ln *k*_cat_ for the two-state model (circles, enzyme colour) and the piecewise reference (squares, gray).

MbPFK/GK does exhibit single-regime behaviour across its entire measured temperature range (10–71 °C). Notably, MbPFK/GK combines high thermal stability with efficient low-temperature catalysis, an observation that does not align with the expectation of reduced stability in psychrophilic enzymes [2], and is consistent with recent systematic analyses questioning the universality of this trend [6].

The ancestral enzyme ancM displayed a two-regime 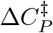 profile similar to that of MmPFK/GK, with transitions occurring at higher temperatures consistent with its greater thermal stability. The enhanced thermostability of ancM relative to the mesophilic descendant MmPFK/GK is consistent with the trend observed in other resurrected ancestral proteins [31,32].

### Domain-closure dynamics as a determinant of the 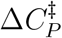 magnitude

The single-regime 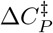 of MbPFK/GK (− 2.65 ± 0.04 kJ mol^−1^ K^−1^) is the most negative among the three enzymes in their respective moderate-temperature regimes, consistent with a greater intrinsic conformational restriction upon transition-state formation. As shown in Figure 4B, the ordering of 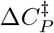 values across MbPFK/GK, M4MbPFK/GK, and MmPFK/GK follows the sequence of domain-closure modes established by molecular dynamics simulations [25]: breathing (−2.65 kJ mol^−1^ K^−1^) > intermediate (−0.99 kJ mol^−1^ K^−1^) > twist (≈ 0). One possibility is that the distinct “breathing-type” domain dynamics of MbPFK/GK [25] preclude the conformational reorganization underlying the second regime. This consistent ordering supports the view that the mode of domain-closure motion determines the degree of conformational restriction imposed on the transition state.

Walker et al. established that 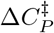 encodes mechanistically meaningful information about the conformational landscape explored during catalysis, arising from cooperative transitions between conformational states of the enzyme– substrate complex that remodel hydrogen-bonding networks and active-site order [17]. This relationship between protein dynamics and 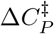 finds independent support in other enzyme systems. For MalL, the distal rigidifying mutation V200S halves the standard MMRT 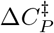 from −11.6 ± 0.4 to −5.9 ± 0.5 kJ mol^−1^ K^−1^ [33], and also reduces the high-temperature plateau in the MMRT-2S model from −28.1 ± 6.3 to −16.3 ± 0.9 kJ mol^−1^ K^−1^ [17]. Similarly, directed evolution and scaffold rigidification of the *de novo* Kemp eliminase induced a negative 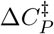 where none existed in the original design, establishing the emergence of 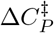 as a hallmark of optimized catalytic efficiency [14,15]. Molecular dynamics simulations have shown that the tightening underlying 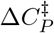 extends beyond the active site to encompass whole-protein dynamical networks [12]. Together with our findings for M4MbPFK/GK, these observations suggest that 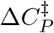 is a sensitive reporter of the global conformational dynamics of the enzyme–substrate complex.

However, the physical basis underlying the exceptionally large 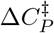 values observed near the thermal stability limits of ancM and MmPFK/GK remains to be elucidated.

## Conclusions

Our comparative analysis of three homologous ADP-dependent PFK/GK enzymes from distinct thermal niches shows that a temperature-dependent 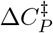 can manifest as two distinct regimes within a single enzyme, providing direct support for the proposal that a temperature-dependent 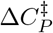 is a general feature of enzyme catalysis [17]. MmPFK/GK and ancM both display two-regime 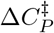 profiles, with a transition to highly negative values near their respective thermal stability limits. The magnitudes in the high-temperature regime (− 44 and − 36 kJ mol^−1^ K^−1^) far exceed previously reported values, and the combined kinetic and CD evidence indicates that they arise from catalytically competent species undergoing pre-melting conformational changes rather than from denaturation. At moderate temperatures, the ordering of 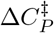 across MbPFK/GK, M4MbPFK/GK, and MmPFK/GK correlates with their mode of domain-closure dynamics, linking the degree of transition-state conformational restriction to a defined structural property. A two-pathway model, in which both the free enzyme and the enzyme–substrate complex populate two catalytically competent conformations in a temperature-dependent equilibrium, can account for the dual-regime behaviour observed across the entire temperature range and provides a plausible mechanistic basis for the abrupt transition to highly negative 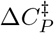 values. Whether this behaviour is a general feature of enzyme catalysis or is confined to structural architectures remains unknown and will require comparative analyses spanning diverse enzyme families.

## Materials and methods

### Protein expression and purification

Enzymes were expressed and purified as previously described in Zamora et al. [25]. Briefly, *E. coli* strain BL21(DE3) was transformed with pET-28b plasmids containing the cDNA of ADP-PFK/GK from *M. burtonii*, ADP-PFK/GK from *M. maripaludis*, ADP-PFK/GK ancM, the glucose-6-phosphate dehydrogenase from *Thermotoga maritima* (TmG6PDH), and the G6PDH from *Leuconostoc mesenteroides* (LmG6PDH). For the ADP-PFK/GK from *M. burtonii*, cells were cultured at 37 °C in LB broth containing 35 *µ*g*·*mL^−1^ kanamycin and grown until OD_600_ of approximately 1.0. For the expressions of ADP-PFK/GK from *M. maripaludis*, ADP-PFK/GK ancM, TmG6PDH, and LmG6PDH, cells were cultured at 37 °C in LB broth containing 35 *µ*g*·*mL^−1^ kanamycin and grown until OD_600_ of 0.4–0.5. Protein expression was induced in all proteins by adding 1 mM IPTG. Then, cultures were incubated at 14 °C overnight for ADP-PFK/GK from *M. burtonii* and at 30 °C for ADP-PFK/GK from *M. maripaludis* and ancM. TmG6PDH and LmG6PDH were expressed at 37 °C for 4 hours. Cells were collected by centrifugation, suspended in binding buffer (25 mM Tris-HCl pH 7.8, 500 mM NaCl, 20 mM imidazole, and 5 mM MgCl_2_), and disrupted by sonication. After centrifugation (18,514 *×* g for 30 min, at 4 °C) the soluble fraction was loaded onto a Ni^2+^-NTA affinity column (HisTrap HP, Cytiva, UK), and protein was eluted with a linear gradient of imidazole between 20 and 500 mM. Fractions with ADP-PFK/GK activity were pooled, concentrated, and dialyzed against 25 mM Tris-HCl pH 7.8, 200 mM NaCl, and 5 mM MgCl_2_ buffer. TmG6PDH and LmG6PDH were dialyzed against 25 mM Tris-HCl pH 7.8 and 200 mM NaCl. All proteins were supplemented with 20% glycerol, flash-frozen in liquid nitrogen, and stored at − 80 °C. Enzyme purity was assessed by SDS-PAGE. Protein concentration was measured using the Bradford assay.

### Enzyme kinetic assays

Kinetic measurements were performed using a JASCO V-730 spectrophotometer coupled with a Peltier PACT-743. The standard assay reaction of glucose saturation curves for ADP-PFK/GK enzymes was carried out in a final volume of 0.5 mL and contained 50 mM HEPES buffer pH 7.0, saturating MgADP concentrations, and 1 mM free Mg^2+^. For temperatures between 10 and 45 °C, LmG6PDH (mesophilic) and 0.2 mM NAD^+^ were used as the auxiliary assay. For temperatures between 50 and 83 °C, TmG6PDH and 0.2 mM NADP^+^ were used. Both auxiliary enzymes were in excess and adequately controlled for not being rate-limiting (LmG6PDH and TmG6PDH at 5 U per assay, Table S4, Table S3). The free Mg^2+^ concentration was calculated from the total concentrations of ADP and Mg^2+^ as described previously [34]. Saturating concentrations of MgADP were determined by estimating the Michaelis–Menten constant (*K*_*m*_) for this substrate at representative low and high temperatures, using 100 mM glucose (Table S2). Concentrations of 5–10 times the ADP – Mg^2+^ *K*_*m*_ for all the enzymes were used to generate glucose saturation curves at different temperatures.

Initial velocities were obtained by measuring absorbance at 340 nm and performing linear regression analysis using the manufacturer’s software (Spectra Manager™ for V-600/700 and FP-8000 series; Jasco, Japan). To calculate the specific activity (U/mg, where 1 U is 1 *µ*mol of product transformed per minute) an extinction coefficient of 6.22 mM^−1^ *·*cm^−1^ was used.

The Michaelis-Menten equation (Eq. 8) and the substrate inhibition model (Eq. 9) were fitted to the initial velocities curves by non-linear regression analysis, using the SciPy library [35].

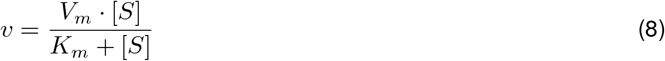

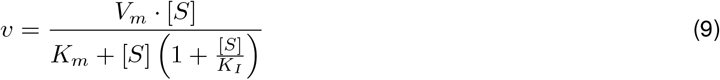

The preferred model was selected based on the Corrected Akaike Information Criterion (AICc) [36].

### Calculation of thermodynamic activation parameters

To estimate 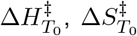, and 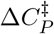, the Eyring equation and the MMRT model (Eq. 1) were fitted to the *k*_cat_ vs. temperature curves by non-linear regression analysis, using the SciPy library [35]. Fits were weighted by the inverse square of the propagated uncertainty on *k*_cat_. The reference temperature (*T*_0_) was set to the midpoint of each fitted temperature range. The fitted 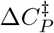 is invariant with respect to the choice of the reference temperature *T*_0_ (Figure S4). Parameter uncertainties were estimated from the covariance matrix of the final fit. For MmPFK/GK and ancM, independent fits were performed for two temperature regimes. The regime boundary was placed at the highest temperature consistent with the low-T fit, identified by iteratively excluding high-temperature data points and monitoring the stability of the fitted 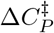. For each regime, both the classical Eyring equation 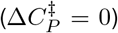 and the full MMRT model were fitted, and the preferred model was selected based on the AIC (Table S5). The extended MMRT-1L (Eq. 4) and MMRT-2S (Eq. 3) models were also fitted over the full temperature range for each enzyme, with model selection by AIC (Table S8).

### Circular dichroism spectroscopy

Far-UV CD spectra were measured using JASCO J-720 and J-815 spectropolarimeters equipped with temperature control. For these measurements, proteins were dialyzed against a buffer containing 6 mM MOPS (pH 7.0) and 15 mM NaCl. Six repeat scans for each sample were measured over the wavelength range from 270 to 200 nm, with a 1 nm interval, in quartz cells of 0.1 cm path length. Six spectra were averaged, baseline spectra subtracted, and the net spectrum was smoothed using a Savitzky-Golay filter using CDToolX [37]. Final processed spectra were expressed in Δ*ε* (M^−1^ *·*cm^−1^) using mean residue weights of 114.1, 111.9 and 115.9 Da, for MmPFK/GK, MbPFK/GK and ancM, respectively. Measurements with an HT value higher than 700 V were not considered.

Quantitative secondary structure fractions were estimated by non-negative least-squares (NNLS) decomposition of the far-UV CD spectra against the SP175 reference dataset [37], which comprises 71 soluble proteins with known crystallographic secondary structure fractions. Each query spectrum was expressed as a non-negative linear combination of the reference protein spectra subject to a soft sum-to-one constraint, simultaneously recovering fractions of α-helix, *β*-sheet, turn, and disordered content. The fitting window extended from the actual lowest reliably measured wavelength of each spectrum (203–208 nm) to 240 nm, the upper limit of the SP175 dataset. Fit quality was assessed by the normalised root-mean-square deviation (NRMSD) between the measured and reconstructed spectra.

### Thermal denaturation

Thermal denaturation was monitored over a temperature range from 25 °C to 70–90 °C by following the ellipticity at 222 nm at a heating rate of 1 °C/min. The fraction unfolded was determined by normalising the observed ellipticity to the difference between the native and fully denatured baselines. Since thermal denaturation of the studied proteins is irreversible, an apparent melting temperature 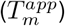 was determined by fitting a phenomenological sigmoidal equation with linear baselines for the native and denatured states (Eq. 10) to the normalised data by non-linear regression analysis in Python using the SciPy library [35]:

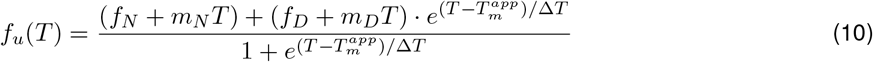

where *f*_*N*_ and *f*_*D*_ are the intercepts of the native and denatured baselines, *m*_*N*_ and *m*_*D*_ are their respective slopes, 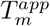 is the apparent melting temperature, and Δ*T* is a parameter describing the width of the transition.

### Unfolding kinetics

Unfolding kinetics were monitored by CD at 222 nm as a function of time. Enzyme solutions in 6 mM MOPS, pH 7.0, and 15 mM NaCl, supplemented with 0, 10, 100, or 500 mM glucose, were placed in the spectropolarimeter holder, which had been previously equilibrated to the target temperature. A first-order exponential model was fitted to the time-dependent CD signal (Eq. 11) by non-linear regression analysis in Python using the SciPy library [35] (nonlinear least-squares):

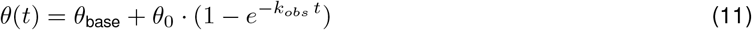

where *θ*(*t*) is the CD signal at 222 nm at time *t, θ*_base_ is the baseline ellipticity, *θ*_0_ is the total amplitude of the signal change, and *k*_*obs*_ is the observed first-order rate constant. For MmPFK/GK and ancM, the initial 30–120 s of data was excluded to avoid thermal equilibration artifacts, with the exclusion window adjusted per temperature (Figure S6).

When the CD trace was nearly linear, indicating very slow unfolding, *k*_*obs*_ was estimated from the slope of a linear fit divided by the fixed *θ*_0_. To assess the kinetic competence of the enzymes relative to their conformational transition rates, the ratio *k*_*cat*_*/k*_*obs*_ was computed at each temperature by using the *k*_cat_ values at the temperatures used for the unfolding kinetics measurements.

## Abbreviations

CD: circular dichroism
GK: glucokinase
MMRT: macromolecular rate theory
PFK: phosphofructokinase
TLC: transition-state-like conformation

## Author contributions

IAV, PM, VG, GVB, VCF and FGO planned experiments; IAV, PM, LHC and FGO performed experiments; IAV, GVB and PM analysed data; VG and VCF contributed reagents or other essential material; IAV, VG and GVB wrote the paper.

## Acknowledgements

The authors thank Dr. Andressa P.A. Pinto and Dr. Richard Garrat from the Biophysics and Structural Biology Group at the Instituto de Física de São Carlos, Universidade de São Paulo, for their generous access to the Jasco J-720 and J-815 spectropolarimeters and to Dr. Richard Garrat for financial support to IAV via the IFSC International Internship Program.

