## Supplementary Information for "Two activation heat capacity regimes underlie temperature-dependent catalysis in homologous archaeal ADP-dependent kinases"

**Index**

- Table S1. Fitted kinetic parameters for all enzymes ........ 2
- Table S2. Fitted parameters from ADP$-$Mg$^{2+}$ saturation curves at representative temperatures ........ 3
- Table S3. TmG6PDH specific activity vs temperature ........ 3
- Table S4. LmG6PDH specific activity ($V_{\mathrm{max}}$) vs temperature at saturating G6P ........ 3
- Figure S1. Temperature dependence of the substrate inhibition constant ($K_I$) ........ 4
- Figure S2. Comparison of Eyring and MMRT fits ........ 5
- Table S5. Comparison of Eyring and MMRT model fits ........ 5
- Figure S3. Thermodynamic decomposition of MMRT activation parameters ........ 6
- Figure S4. Dependence of MMRT activation parameters on $T_0$ ........ 6
- Figure S5. LmG6PDH kinetic characterization ........ 7
- Table S6. Comparison of Eyring and MMRT model fits for LmG6PDH ........ 7
- Figure S6. CD-monitored thermal denaturation kinetics ........ 8
- Figure S7. $k_{\mathrm{obs}}$ vs temperature: unfolding rate constants and exponential fits ........ 9
- Figure S8. M4MbPFK/GK kinetic characterization ........ 10
- Table S7. Comparison of Eyring and MMRT model fits for M4MbPFK/GK ........ 10
- Figure S9. Global model fits: Eyring, MMRT, MMRT-1L, and MMRT-2S ........ 11
- Table S8. AICc model comparison: Eyring vs MMRT vs MMRT-1L vs MMRT-2S ........ 12

Table S 1: Kinetic parameters from glucose saturation curves at each temperature, from fits to the Michaelis–Menten equation (MbPFK/GK, ancM) or the substrate inhibition model (MmPFK/GK; Eq. 9). Mean $\pm$ standard error.

| **MmPFK/GK** | | | | **MbPFK/GK** | | | **ancM** | | |
|---|---|---|---|---|---|---|---|---|---|
| $T$ (°C) | $k_{\text{cat}}$ (s$^{-1}$) | $K_m$ (mM) | $K_I$ (mM) | $T$ (°C) | $k_{\text{cat}}$ (s$^{-1}$) | $K_m$ (mM) | $T$ (°C) | $k_{\text{cat}}$ (s$^{-1}$) | $K_m$ (mM) |
| 10.5 | $11.2 \pm 0.6$ | $57.3 \pm 5.9$ | $1711 \pm 375$ | 10.0 | $0.44 \pm 0.02$ | $4.2 \pm 0.6$ | 25.0 | $0.90 \pm 0.02$ | $6.5 \pm 0.7$ |
| 15.0 | $22.3 \pm 1.0$ | $44.0 \pm 4.0$ | $605 \pm 71$ | 15.0 | $0.78 \pm 0.01$ | $4.2 \pm 0.3$ | 30.0 | $1.12 \pm 0.02$ | $6.0 \pm 0.5$ |
| 19.8 | $44.5 \pm 4.4$ | $43.2 \pm 7.7$ | $341 \pm 69$ | 20.0 | $1.41 \pm 0.02$ | $5.0 \pm 0.2$ | 35.0 | $1.26 \pm 0.03$ | $6.1 \pm 0.6$ |
| 25.1 | $56.8 \pm 3.1$ | $20.9 \pm 2.3$ | $344 \pm 46$ | 25.0 | $3.21 \pm 0.07$ | $5.7 \pm 0.5$ | 40.0 | $1.46 \pm 0.05$ | $8.0 \pm 1.1$ |
| 30.4 | $86.6 \pm 8.5$ | $27.1 \pm 4.9$ | $265 \pm 55$ | 30.0 | $7.56 \pm 0.26$ | $8.0 \pm 0.9$ | 45.0 | $1.73 \pm 0.05$ | $7.0 \pm 0.9$ |
| 35.2 | $97.2 \pm 5.8$ | $23.7 \pm 2.9$ | $450 \pm 69$ | 35.0 | $10.6 \pm 0.1$ | $9.6 \pm 0.4$ | 50.0 | $1.94 \pm 0.05$ | $7.1 \pm 0.9$ |
| 40.3 | $133.9 \pm 7.6$ | $6.2 \pm 1.1$ | $753 \pm 171$ | 40.0 | $12.1 \pm 0.3$ | $18.0 \pm 1.3$ | 54.5 | $2.29 \pm 0.04$ | $6.5 \pm 0.5$ |
| 45.6 | $190.7 \pm 13.5$ | $4.0 \pm 1.0$ | $808 \pm 254$ | 45.0 | $14.7 \pm 0.3$ | $25.5 \pm 1.9$ | 60.0 | $2.59 \pm 0.03$ | $7.0 \pm 0.4$ |
| 50.9 | $286.8 \pm 10.4$ | $3.5 \pm 0.4$ | $882 \pm 151$ | 50.0 | $18.0 \pm 1.9$ | $23.6 \pm 8.3$ | 66.5 | $2.83 \pm 0.14$ | $8.9 \pm 1.7$ |
| 55.7 | $446.5 \pm 7.0$ | $4.0 \pm 0.2$ | $702 \pm 46$ | 55.0 | $19.6 \pm 0.5$ | $37.8 \pm 2.7$ | 73.8 | $3.41 \pm 0.11$ | $7.0 \pm 1.0$ |
| 56.8 | $635.5 \pm 50.9$ | $9.1 \pm 2.1$ | $805 \pm 251$ | 58.5 | $21.3 \pm 1.4$ | $45.9 \pm 11.5$ | 75.5 | $4.41 \pm 0.13$ | $10.5 \pm 1.2$ |
| 57.8 | $808 \pm 121$ | $21.8 \pm 8.6$ | $1901 \pm 1536$ | 63.6 | $21.9 \pm 1.2$ | $36.0 \pm 8.3$ | 76.3 | $4.30 \pm 0.34$ | $6.1 \pm 2.2$ |
| 60.2 | $948.5 \pm 34.1$ | $18.5 \pm 1.7$ | $1008 \pm 137$ | 65.5 | $25.4 \pm 0.9$ | $114.7 \pm 12.0$ | 76.3 | $4.72 \pm 0.10$ | $7.4 \pm 0.7$ |
| 61.8 | $932.7 \pm 45.4$ | $18.6 \pm 2.4$ | $1195 \pm 242$ | 66.6 | $25.2 \pm 1.1$ | $107.0 \pm 13.3$ | 76.6 | $4.92 \pm 0.08$ | $7.8 \pm 0.5$ |
| 62.9 | $781.4 \pm 47.3$ | $17.1 \pm 2.9$ | $2126 \pm 793$ | 67.2 | $22.4 \pm 2.1$ | $120.6 \pm 33.3$ | 77.5 | $5.80 \pm 0.16$ | $8.9 \pm 1.0$ |
| 66.0 | $587.9 \pm 40.2$ | $16.8 \pm 3.1$ | $1313 \pm 404$ | 68.7 | $20.8 \pm 1.3$ | $136.6 \pm 22.6$ | 78.3 | $6.22 \pm 0.11$ | $7.9 \pm 0.6$ |
| | | | | 69.5 | $23.1 \pm 3.8$ | $134.9 \pm 59.0$ | 78.9 | $5.95 \pm 0.42$ | $13.9 \pm 5.1$ |
| | | | | 70.7 | $19.0 \pm 1.0$ | $121.0 \pm 18.5$ | 79.3 | $6.75 \pm 0.14$ | $7.7 \pm 0.6$ |
| | | | | | | | 80.1 | $6.31 \pm 0.10$ | $13.3 \pm 4.1$ |
| | | | | | | | 81.5 | $5.86 \pm 0.34$ | $6.8 \pm 14.8$ |
| | | | | | | | 82.1 | $6.29 \pm 0.14$ | $8.1 \pm 5.5$ |
| | | | | | | | 83.0 | $5.45 \pm 0.07$ | $7.8 \pm 3.5$ |

Table S 2: ADP$-$Mg$^{2+}$ kinetic parameters at representative temperatures, used to set the saturating ADP$-$Mg$^{2+}$ concentration ($\geq 10 \times K_m$) for the glucose saturation curves (Table S1). Michaelis–Menten fits; mean $\pm$ standard error.

| **MmPFK/GK** | | | **MbPFK/GK** | | | **ancM** | | |
|---|---|---|---|---|---|---|---|---|
| $T$ (℃) | $k_{cat}$ (s$^{-1}$) | $K_m$ (mM) | $T$ (℃) | $k_{cat}$ (s$^{-1}$) | $K_m$ (mM) | $T$ (℃) | $k_{cat}$ (s$^{-1}$) | $K_m$ (mM) |
| 25 | $44.9 \pm 1.5$ | $0.12 \pm 0.02$ | 25 | $2.24 \pm 0.08$ | $0.19 \pm 0.02$ | 25 | $0.97 \pm 0.06$ | $0.18 \pm 0.04$ |
| 45 | $169.1 \pm 2.6$ | $0.19 \pm 0.01$ | 45 | $10.96 \pm 0.61$ | $0.31 \pm 0.06$ | 45 | $1.90 \pm 0.11$ | $0.19 \pm 0.04$ |
| 55 | $446.6 \pm 13.5$ | $0.28 \pm 0.03$ | – | – | – | 55 | $2.25 \pm 0.07$ | $0.22 \pm 0.02$ |
| 65 | $549.1 \pm 25.4$ | $0.98 \pm 0.12$ | 65.1 | $23.5 \pm 3.4$ | $4.15 \pm 1.74$ | – | – | – |

Table S 3: Specific activity of TmG6PDH vs temperature. Mean $\pm$ standard deviation of measurements at 10, 50, and 100 nM TmG6PDH.

| $T$ (℃) | U/mg ($n = 3$) |
|---|---|
| 35.0 | $2.08 \pm 0.09$ |
| 40.0 | $3.36 \pm 0.15$ |
| 45.0 | $5.66 \pm 0.18$ |
| 50.2 | $7.89 \pm 0.23$ |
| 55.1 | $9.86 \pm 0.26$ |
| 60.0 | $12.25 \pm 1.31$ |
| 64.8 | $16.91 \pm 1.39$ |
| 69.6 | $22.54 \pm 1.60$ |
| 74.4 | $28.35 \pm 2.52$ |
| 79.9 | $35.50 \pm 4.46$ |
| 84.5 | $49.18 \pm 2.62$ |

Table S 4: Specific activity ($V_{max}$, U/mg) of LmG6PDH at saturating G6P vs temperature, from the fits in Figure S5. Mean $\pm$ standard error.

| $T$ (℃) | $V_{max}$ (U/mg) |
|---|---|
| 10.0 | $144.9 \pm 6.4$ |
| 15.0 | $176.3 \pm 3.4$ |
| 20.2 | $230.5 \pm 14.6$ |
| 25.0 | $311.9 \pm 8.0$ |
| 30.0 | $352.8 \pm 13.8$ |
| 35.0 | $469.0 \pm 12.6$ |
| 40.0 | $598.5 \pm 21.9$ |
| 43.8 | $703.9 \pm 26.4$ |
| 48.0 | $827.0 \pm 65.8$ |

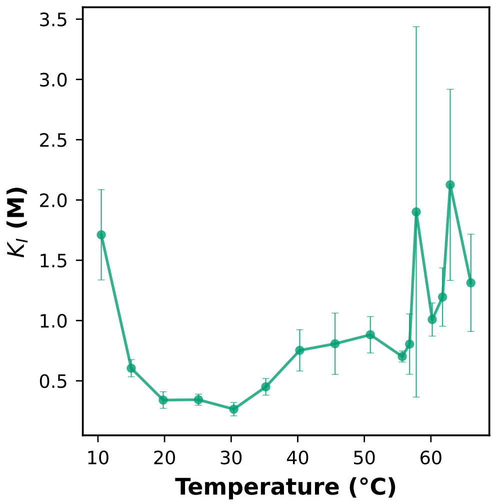


Figure S 1: Temperature dependence of the substrate inhibition constant ($K_I$) for MmPFK/GK, from fits to the substrate inhibition model (Eq. 9). $K_I$ stays near 0.3–0.9 M between 20 and 55 ℃ and rises sharply near $T_m^{app}$. Error bars, standard error.

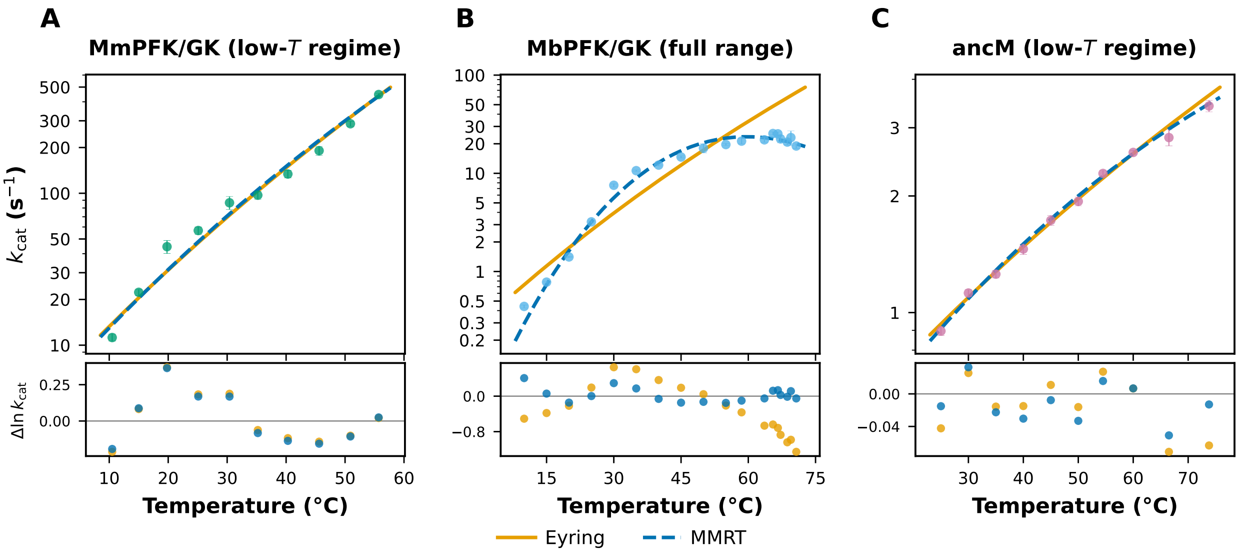


Figure S 2: Eyring vs MMRT fits to $k_{\text{cat}}(T)$ (top) with residuals (bottom): Eyring (orange, solid), MMRT (blue, dashed). **(A)** MmPFK/GK low-$T$ regime (10.5–55.7 ℃). **(B)** MbPFK/GK full range (10.0–70.7 ℃). **(C)** ancM low-$T$ regime (25.0–73.8 ℃). Statistics in Table S5.

Table S 5: Eyring vs MMRT fit parameters (low-$T$ regime for MmPFK/GK and ancM). $T_0$ is the midpoint of each fitted range; the preferred model (•) was selected by AIC.

| | MmPFK/GK | | MbPFK/GK | | ancM | |
|---|---|---|---|---|---|---|
| Parameter | Eyring | MMRT | Eyring | MMRT | Eyring | MMRT |
| Range (℃) | 10.5–55.7 | | 10.0–70.7 | | 25.0–73.8 | |
| $n$ | 10 | | 18 | | 10 | |
| $T_0$ (℃) | – | 33 | – | 40 | – | 49 |
| $\Delta H^{\ddagger}_{T_0}$ (kJ mol$^{-1}$) | $56.7 \pm 0.6$ | $56.8 \pm 0.7$ | $57.5 \pm 0.3$ | $29.5 \pm 0.5$ | $21.3 \pm 0.4$ | $20.9 \pm 0.5$ |
| $T_0 \Delta S^{\ddagger}_{T_0}$ (kJ mol$^{-1}$) | $-7.0 \pm 0.6$ | $-6.8 \pm 0.7$ | $-14.1 \pm 0.3$ | $-41.4 \pm 0.5$ | $-55.9 \pm 0.4$ | $-56.2 \pm 0.5$ |
| $\Delta G^{\ddagger}_{T_0}$ (kJ mol$^{-1}$) | $63.7 \pm 0.04$ | $63.6 \pm 0.1$ | $71.6 \pm 0.04$ | $70.9 \pm 0.04$ | $77.2 \pm 0.03$ | $77.1 \pm 0.03$ |
| $\Delta C^{\ddagger}_{P}$ (kJ mol$^{-1}$ K$^{-1}$) | 0 | $-0.10 \pm 0.12$ | 0 | $-2.65 \pm 0.04$ | 0 | $-0.17 \pm 0.07$ |
| RMSE | 0.175 | 0.172 | 0.640 | 0.155 | 0.036 | 0.026 |
| AIC | $-30.9$ | $-29.2$ | $-12.1$ | $-61.1$ | $-62.3$ | $-66.7$ |
| Preferred | • | | | • | | • |

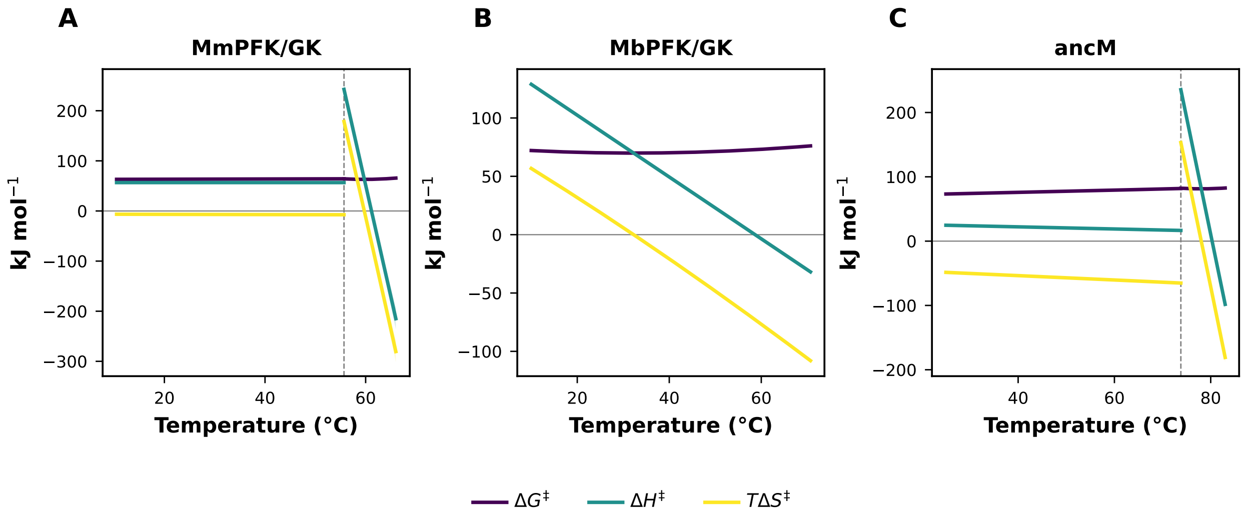


Figure S 3: Temperature dependence of $\Delta G^{\ddagger}$ (purple), $\Delta H^{\ddagger}$ (teal), and $T\Delta S^{\ddagger}$ (yellow) from the MMRT fits (Table 1) for **(A)** MmPFK/GK, **(B)** MbPFK/GK, and **(C)** ancM. Bands, propagated $1\sigma$ uncertainty; dashed lines (A, C) mark the regime boundary.

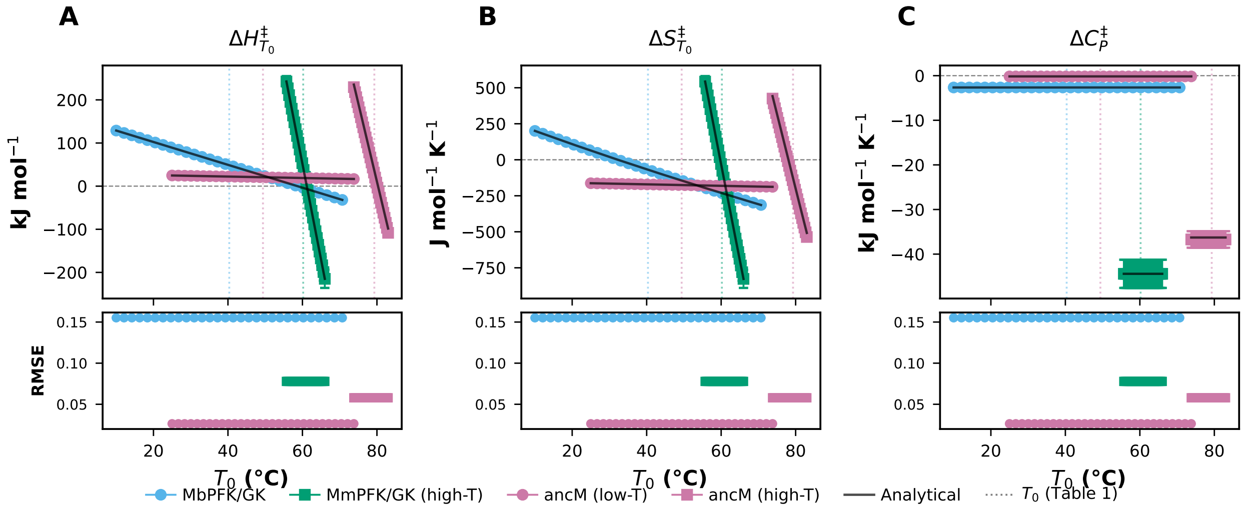


Figure S 4: MMRT activation parameters vs the fixed reference temperature $T_0$: **(A)** $\Delta H^{\ddagger}_{T_0}$, **(B)** $\Delta S^{\ddagger}_{T_0}$, **(C)** $\Delta C^{\ddagger}_{P}$. Black lines, analytical extrapolation from the reference fit; bottom panels, RMSE. Error bars, $1\sigma$; dotted lines mark the $T_0$ values of Table 1. MmPFK/GK low-$T$ (Eyring) is omitted.

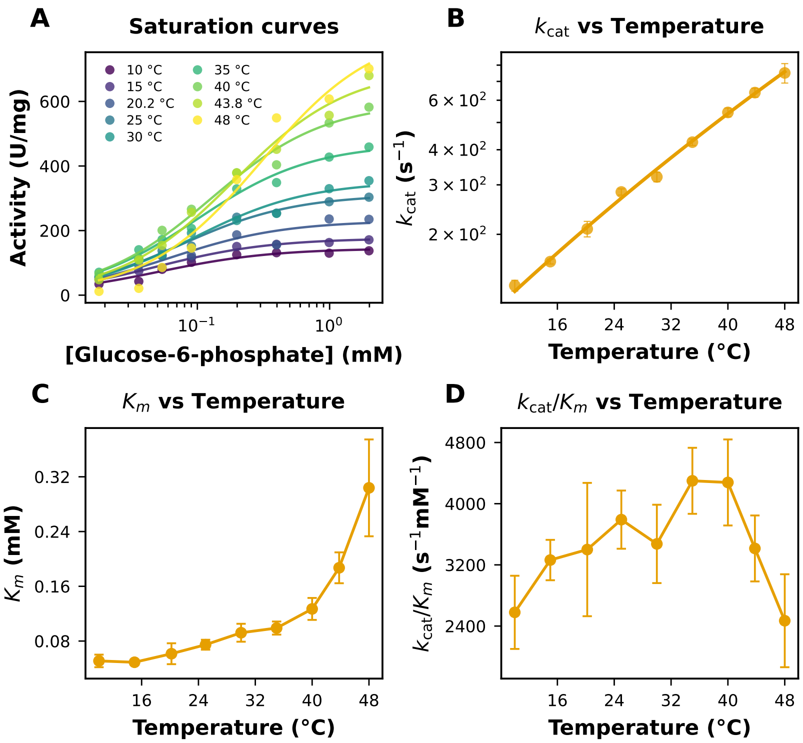


Figure S 5: Kinetic characterisation of the coupling enzyme LmG6PDH. **(A)** Glucose-6-phosphate saturation curves (purple = 10 ℃ to yellow = 48 ℃), fitted to Michaelis–Menten (Eq. 8). **(B)** $k_{\text{cat}}$ vs temperature with Eyring fit. **(C)** $K_m$ and **(D)** $k_{\text{cat}}/K_m$ vs temperature. Mean $\pm$ SEM.

Table S 6: Eyring vs MMRT fit parameters for LmG6PDH. $T_0$ is the midpoint of the fitted range; the preferred model (●) was selected by AIC.

| Parameter | Eyring | MMRT |
|---|---|---|
| Range (℃) | 10.0–48.0 | |
| $n$ | 9 | |
| $T_0$ (℃) | – | 29.0 |
| $\Delta H^{\ddagger}_{T_0}$ (kJ mol$^{-1}$) | $33.3 \pm 0.7$ | $33.3 \pm 0.8$ |
| $T_0 \Delta S^{\ddagger}_{T_0}$ (kJ mol$^{-1}$) | $-26.3 \pm 0.7$ | $-26.3 \pm 0.8$ |
| $\Delta C^{\ddagger}_P$ (kJ mol$^{-1}$ K$^{-1}$) | 0 | $0.006 \pm 0.163$ |
| RMSE | 0.033 | 0.033 |
| AIC | $-57.3$ | $-55.3$ |
| Preferred | ● | |

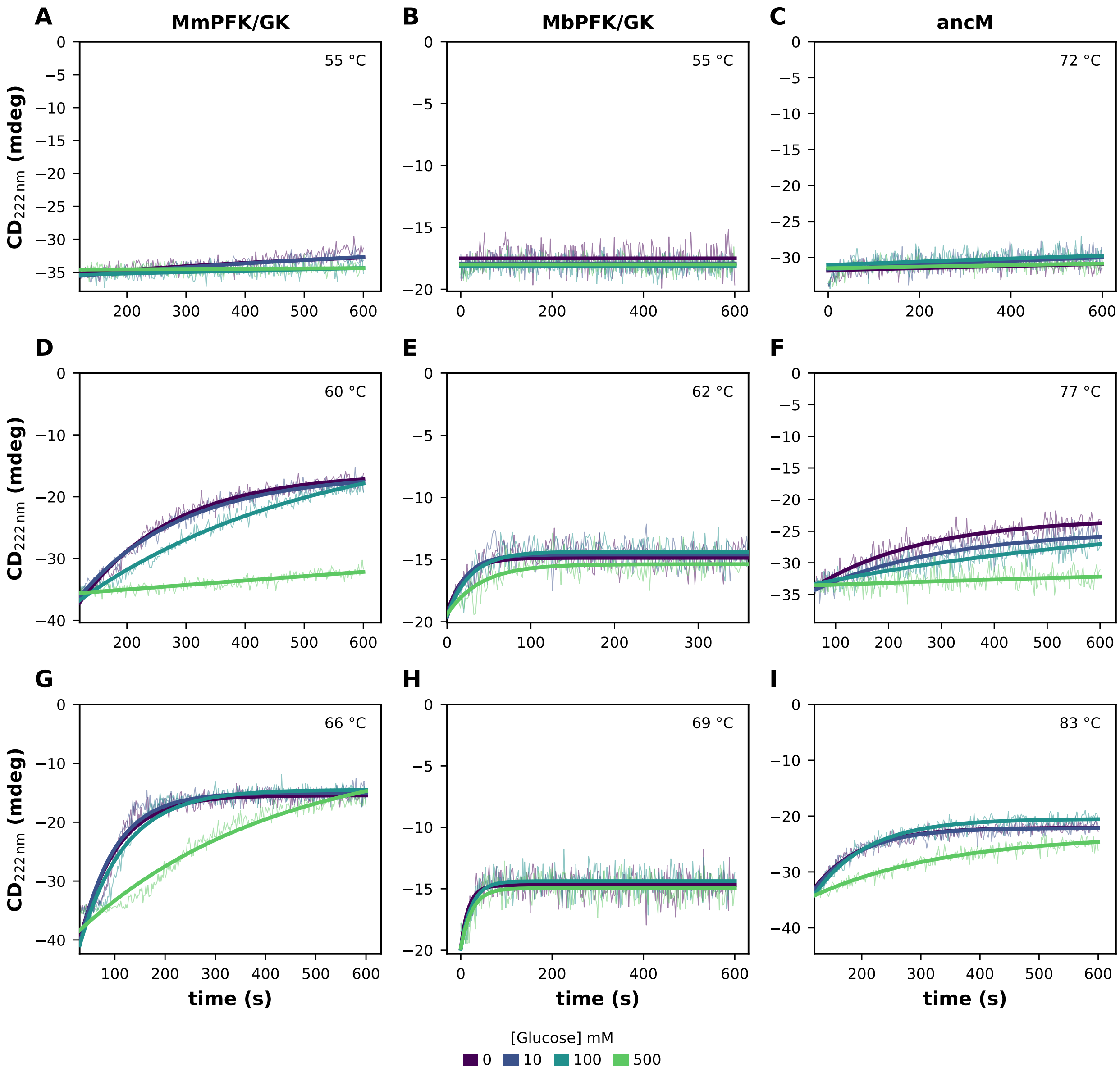


Figure S 6: Time-resolved CD unfolding kinetics (ellipticity at 222 nm) for MmPFK/GK (**A, D, G**), ancM (**B, E, H**), and MbPFK/GK (**C, F, I**) at 0, 10, 100, and 500 mM glucose (colour-coded). Thin lines, raw data; thick lines, fits to Eq. 11. Temperature increases top to bottom within each column.

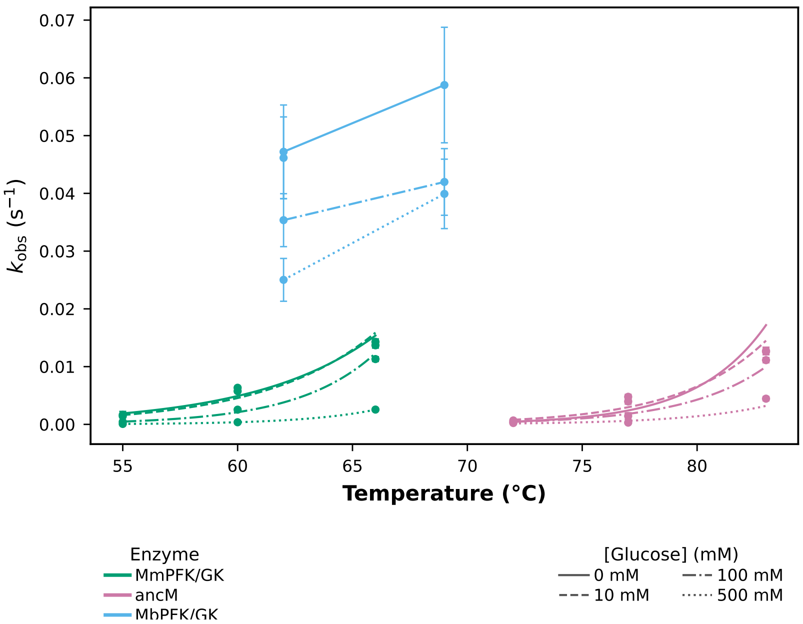


Figure S 7: Observed unfolding rate constant $k_{\mathrm{obs}}$ vs temperature for MmPFK/GK (green), MbPFK/GK (blue), and ancM (pink); line style encodes glucose concentration (legend). Curves are exponential fits ($k_{\mathrm{obs}} = a\, e^{bT}$) for MmPFK/GK and ancM; MbPFK/GK has too few points to fit. MbPFK/GK at 55 ℃ excluded (no detectable CD change). Mean $\pm$ SE.

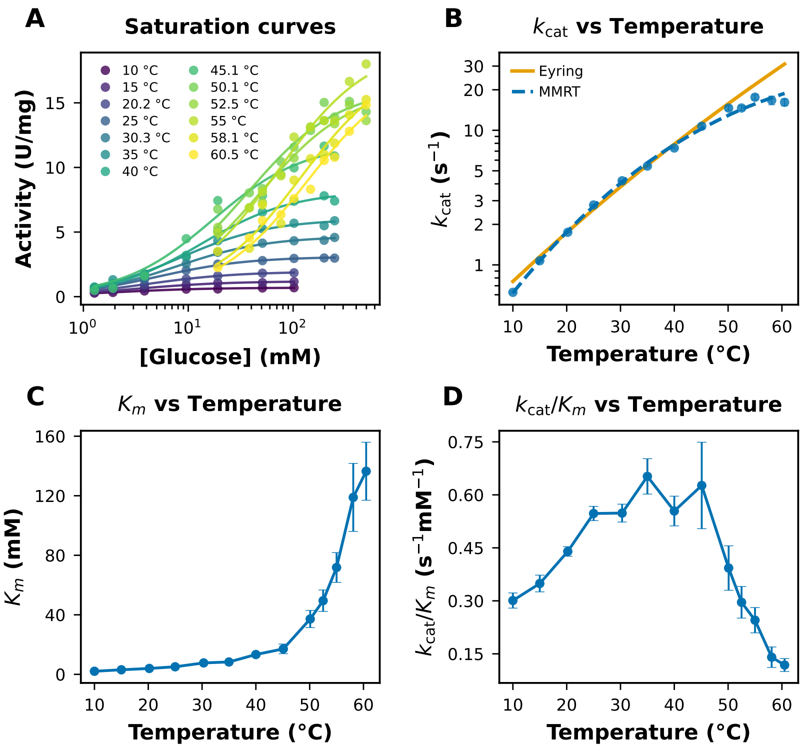


Figure S 8: Kinetic characterisation of M4MbPFK/GK. **(A)** Glucose saturation curves (purple = lowest to yellow = highest temperature), fitted to Michaelis–Menten (Eq. 8). **(B)** $k_{\mathrm{cat}}$ vs temperature with Eyring (solid) and MMRT (dashed) fits. **(C)** $K_m$ and **(D)** $k_{\mathrm{cat}}/K_m$ vs temperature. Mean $\pm$ SEM.

Table S 7: Eyring vs MMRT fit parameters for M4MbPFK/GK. $T_0$ is the midpoint of the fitted range; the preferred model (●) was selected by AIC.

| Parameter | Eyring | MMRT |
|---|---|---|
| Range (℃) | 10.0–60.5 | |
| $n$ | 13 | |
| $T_0$ (℃) | – | 35 |
| $\Delta H^{\ddagger}_{T_0}$ (kJ mol$^{-1}$) | $55.3 \pm 2.4$ | $46.2 \pm 1.4$ |
| $T_0 \Delta S^{\ddagger}_{T_0}$ (kJ mol$^{-1}$) | $-16.1 \pm 2.5$ | $-25.1 \pm 1.4$ |
| $\Delta C^{\ddagger}_{P}$ (kJ mol$^{-1}$ K$^{-1}$) | 0 | $-0.99 \pm 0.11$ |
| RMSE | 0.251 | 0.069 |
| AIC | $-31.9$ | $-63.3$ |
| Preferred | | ● |

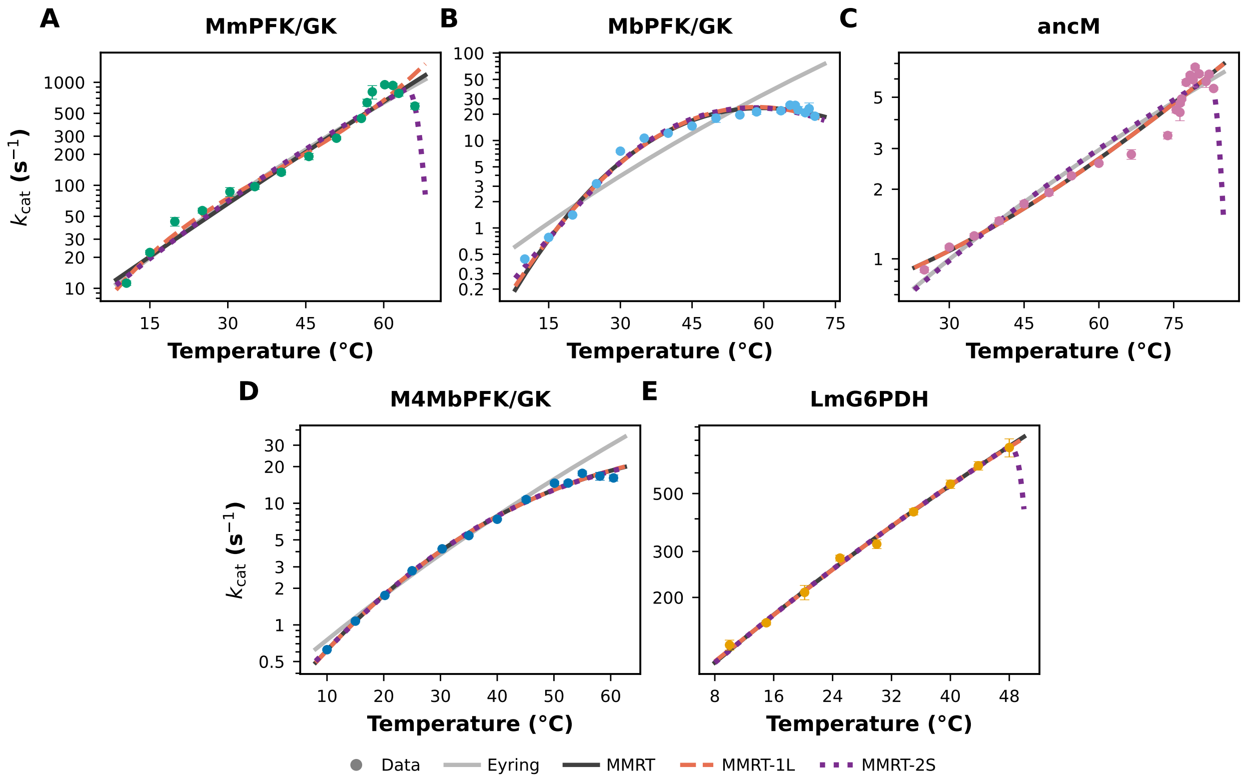


Figure S 9: Global fits of four models (Eyring, MMRT, MMRT-1L, MMRT-2S; 2–5 parameters) to $k_{\mathrm{cat}}(T)$ over the full temperature range of each enzyme. Points, data (enzyme colours); lines, fits (shared legend). MMRT-1L and MMRT-2S traces largely overlap MMRT. AICc comparison in Table S8.

Table S 8: AICc comparison across enzymes (full temperature range). $k$, number of parameters; $T_0$ fixed to the midpoint of each range. Preferred model in bold (lowest AICc); ΔAICc $> 2$ indicates the extra parameters are not justified. The main analysis instead applies Eyring or MMRT piecewise within temperature regimes.

| Enzyme | $n$ | Range (℃) | Model | $k$ | RMSE | AICc | ΔAICc | Preferred |
|---|---|---|---|---|---|---|---|---|
| MmPFK/GK | 16 | 10.5–66.0 | **Eyring** | 2 | 0.251 | $-39.3$ | 0.0 | • |
| | | | MMRT | 3 | 0.257 | $-35.5$ | 3.9 | |
| | | | MMRT-1L | 4 | 0.255 | $-32.0$ | 7.3 | |
| | | | MMRT-2S | 5 | 0.218 | $-32.7$ | 6.6 | |
| MbPFK/GK | 18 | 10.0–70.7 | Eyring | 2 | 0.640 | $-11.3$ | 48.1 | |
| | | | **MMRT** | 3 | 0.155 | $-59.4$ | 0.0 | • |
| | | | MMRT-1L | 4 | 0.158 | $-55.3$ | 4.1 | |
| | | | MMRT-2S | 5 | 0.145 | $-54.6$ | 4.7 | |
| ancM | 22 | 25.0–83.0 | Eyring | 2 | 0.130 | $-85.1$ | 4.5 | |
| | | | **MMRT** | 3 | 0.110 | $-89.6$ | 0.0 | • |
| | | | MMRT-1L | 4 | 0.110 | $-86.6$ | 3.0 | |
| | | | MMRT-2S | 5 | 0.130 | $-75.9$ | 13.8 | |
| LmG6PDH | 9 | 10.0–48.0 | **Eyring** | 2 | 0.033 | $-55.3$ | 0.0 | • |
| | | | MMRT | 3 | 0.033 | $-50.5$ | 4.7 | |
| | | | MMRT-1L | 4 | 0.032 | $-43.7$ | 11.6 | |
| | | | MMRT-2S | 5 | 0.033 | $-31.3$ | 23.9 | |
| M4MbPFK/GK | 13 | 10.0–60.5 | Eyring | 2 | 0.251 | $-30.7$ | 30.0 | |
| | | | **MMRT** | 3 | 0.069 | $-60.7$ | 0.0 | • |
| | | | MMRT-1L | 4 | 0.070 | $-56.0$ | 4.7 | |
| | | | MMRT-2S | 5 | 0.068 | $-51.1$ | 9.5 | |
